# An Automated SNP-Based Approach for Contaminant Identification in Biparental Polyploid Populations of Tropical Forage Grasses

**DOI:** 10.1101/2021.07.01.450796

**Authors:** Felipe Bitencourt Martins, Aline da Costa Lima Moraes, Alexandre Hild Aono, Rebecca Caroline Ulbricht Ferreira, Lucimara Chiari, Rosangela Maria Simeão, Sanzio Carvalho Lima Barrios, Mateus Figueiredo Santos, Liana Jank, Cacilda Borges do Valle, Bianca Baccili Zanotto Vigna, Anete Pereira de Souza

**Affiliations:** Center for Molecular Biology and Genetic Engineering (CBMEG), University of Campinas (UNICAMP), Campinas, São Paulo, Brazil; Embrapa Gado de Corte, Brazilian Agricultural Research Corporation, Campo Grande, Mato Grosso do Sul, Brazil; Embrapa Pecuária Sudeste, Brazilian Agricultural Research Corporation, São Carlos, São Paulo, Brazil; Department of Plant Biology, Biology Institute, University of Campinas (UNICAMP), Campinas, São Paulo, Brazil

**Keywords:** GBS, apomictic clones, self-fertilization, half-siblings, clustering analysis, principal component analysis, allele dosage, Shiny

## Abstract

Artificial hybridization plays a fundamental role in plant breeding programs since it generates new genotypic combinations that can result in desirable phenotypes. Depending on the species and mode of reproduction, controlled crosses may be challenging, and contaminating individuals can be introduced accidentally. In this context, the identification of such contaminants is important to avoid compromising further selection cycles, as well as genetic and genomic studies. The main objective of this work was to propose an automated multivariate methodology for the detection and classification of putative contaminants, including apomictic clones, self-fertilized individuals, half-siblings and full contaminants, in biparental polyploid progenies of tropical forage grasses. We established a pipeline to identify contaminants in genotyping-by-sequencing (GBS) data encoded as allele dosages of single nucleotide polymorphism (SNP) markers by integrating principal component analysis (PCA), genotypic analysis (GA) measures based on Mendelian segregation and clustering analysis (CA). The combination of these methods allowed the correct identification of all contaminants in all simulated progenies and the detection of putative contaminants in three real progenies of tropical forage grasses, providing an easy and promising methodology for the identification of contaminants in biparental progenies of tetraploid and hexaploid species. The proposed pipeline was made available through the polyCID Shiny app and can be easily coupled with traditional genetic approaches, such as linkage map construction, thereby increasing the efficiency of breeding programs.

## 1 Introduction

The concept of artificial crossings to generate experimental plant populations was introduced scientifically in Mendel’s historical work (Mendel, 1866) and became a fundamental tool for genetics studies and breeding programs, maximizing genetic gains by the selection of superior genotypes (Bourke et al., 2018). Although this concept is well known and applied in important crops (Goulet et al., 2017), there are few commercial cultivars of tropical forage grasses originating from artificial hybridization (Azevedo et al., 2019). Perennial tropical forage grasses are recognized worldwide for their economic importance as food for beef and dairy cattle in tropical and subtropical regions (Pereira et al., 2018a; ABIEC, 2020). In addition to recently initiated breeding programs and long selection cycles, some intrinsic biological characteristics, including different reproductive modes (sexual and facultative apomixis), levels of ploidy and self-incompatibility (SI) within and between species, are challenges faced by breeders when performing controlled crosses using these plants (Lutts et al., 1991; Jank et al., 2011; Pereira et al., 2018a; Worthington et al., 2019).

Apomixis is a type of asexual reproduction through seeds that produces progeny genetically identical to the maternal plant (Bicknell, 2004; Hand and Koltunow, 2014). Thus, to explore the genetic diversity of apomictic forage grasses, controlled crosses are performed between sexual and apomictic (pollen donor) parents with contrasting traits and the same ploidy level, which requires previous artificial chromosome duplication of the sexual parent in most species (Pinheiro et al., 2000; Simioni and Valle, 2009; Acuña et al., 2019). However, because of the reproductive system of these plants, during the crosses, some individuals are also generated by foreign pollen or by self-fertilization of female parents. Some of these scenarios can also occur in other species, such as sugarcane, eucalyptus, sainfoin and lettuce (Santos et al., 2014; Subashini et al., 2014; Kempf et al., 2015; Patella et al., 2019). Also, if facultative apomictic plants are used as females, they simultaneously generate hybrid by crossings and clones by apomixis (Smith, 1972). Contamination by physical admixture during seed harvesting and handling is also possible, especially when crosses are performed in the field, as these species are anemophilous (Bateman, 1947; Simeão et al., 2016a). In this context, it is evident that controlled crosses may not avoid contamination, compromising the attainment of pure hybrid progeny and, consecutively, unbiased genetic and genomic methods, such as segregation tests, linkage map construction, quantitative trait locus (QTL) mapping and linkage disequilibrium analysis, which are fundamental for understanding the genotype and its relationship to the phenotype (Kemble et al., 2019).

Traditionally, hybrid identification has been performed on the basis of morphological traits and microsatellite markers (Santos et al., 2014; Jha et al., 2016; Zhao et al., 2017; Patella et al., 2019). However, both methodologies have disadvantages. Morphological traits are time-consuming and have low throughput, with accuracies influenced by environmental factors (Zhao et al., 2017), while developing microsatellite markers is an expensive and time-consuming process that requires previously obtained genomic sequence information and investment in terms of designing locus-specific primers and optimizing polymerase chain reaction (PCR) conditions (Vieira et al., 2016). Moreover, size estimates across alleles at each locus are imprecise, especially in polyploids such as tropical forage grasses, leading to frequent genotyping errors (Guichoux et al., 2011; Hodel et al., 2016). Therefore, there is a need to develop alternative methodologies using molecular markers to quickly and efficiently distinguish true hybrids resulting from planned outcrosses from those resulting from accidental selfing or contamination in biparental populations.

Single nucleotide polymorphism (SNP) markers have been shown to be an excellent tool for genomic studies as a function of their high-throughput nature, low error rates and abundance in eukaryote genomes (Helyar et al., 2011). Additionally, genotyping methodologies based on next-generation sequencing (NGS), such as genotyping-by-sequencing (GBS) proposed by Elshire et al. (2011) and Poland et al. (2012), have been demonstrated to be quick, affordable and highly robust for discovering and profiling a large number of SNP loci, even in species with no available genomic information and large genomes, such as polyploids (Elshire et al., 2011; Poland et al., 2012; Ferreira et al., 2019; Deo et al., 2020; Mollinari et al., 2020). In the last few years, many studies using SNP markers in tropical forage grasses, mainly coupled with principal component analysis (PCA) to investigate the structure of the progenies and remove putative contaminants, have been published (Lara et al., 2019; Deo et al., 2020; Zhang et al., 2020). Even though PCA can be used to retain and explore most of the variation in large SNP datasets through the first principal components (PCs) (Jolliffe and Cadima, 2016), such a multivariate technique is not appropriate for contaminant identification, which requires more specific approaches, such as pedigree reconstruction and sibship and parentage assignment.

The different methods for identifying the parents of a progeny are based on exclusion (Zwart et al., 2016; McClure et al., 2018), likelihood-based (Spielmann et al., 2015) and Bayesian (Christie et al., 2013) techniques, using Mendel’s laws to infer relationships between samples through genotyped loci (Thompson, 1975; Thompson and Meagher, 1987). This evaluation is generally based on pairwise Mendelian segregation tests, comparing individuals and generating different measures that account for the similarity between a sample and one of the parents or for a rate of unexpected genotypes in each sample considering the genotypes of both parents. Therefore, such genotype analyses (GAs) can be used to define what is not genotypically similar and consecutively represents an experimental contaminant. In this work, we propose to use GA measures for performing clustering analyses (CAs) and automatically identifying contaminants in forage grass biparental populations, grouping individuals based on GA similarity measures instead of their raw SNP data. Although CA of large SNP datasets has been extensively used to discover patterns in population relatedness and structure (Gori et al., 2016; Muniz et al., 2019; Yousefi-Mashouf et al., 2021), its use for parentage assignment is not common because of the clusters’ nonspecificity and constancy, but has already been combined with previously described techniques for parentage and sibship inference in diploids (Ellis et al., 2018).

Instead of relying strictly on PCA for population analyses and ad hoc decisions (Deo et al., 2020; Zhang et al., 2020), we created an automated pipeline, combining GA and CA and allowing us not only to precisely identify but also to list the types of contaminants in a biparental cross. For this purpose, we simulated several biparental progenies with contaminants to (1) identify dispersion patterns in a PCA biplot that can suggest the presence of contaminants, (2) create appropriate GA measures for contaminant identification in polyploid forage grass samples, generating scores for all individuals, and (3) integrate such scores in an automatic CA to separate the real hybrids from contaminants. These steps led to the formulation of a unified methodology, which we applied to biparental progenies of three different species of tropical forage grasses: *Megathyrsus maximus* (Jacq.) B. K. Simon & S. W. L. Jacobs (syn. *Panicum maximum* Jacq.), *Urochloa decumbens* (Stapf) R. D. Webster (syn. *Brachiaria decumbens* Stapf) and *Urochloa humidicola* (Rendle) Morrone & Zuloaga (syn. *Brachiaria humidicola* (Rendle) Schweick) (Morrone and Zuloaga, 1992; Torres-González and Morton, 2005). The implemented pipeline was made available through a Shiny app and has a high potential to be employed in prebreeding stages, as well in genomic studies involving polyploid biparental progenies in general.

## 2 Materials and Methods

The following sections describe the steps involved in the generation of real and simulated data and their use to propose a methodology for contaminant identification in biparental crosses. First, the genotyping and allele dosage estimation for biparental F_1_ populations of three tropical forage species is presented (2.1, 2.2 and 2.3). Then, different biparental crossings are simulated (2.4). Finally, contaminant identification methodologies are applied to the simulated and real data (2.5, 2.6, 2.7 and 2.8).

### 2.1 Plant Material

Genotypic data were obtained from biparental F_1_ progenies of *U. humidicola* (2n=6x=36), *U. decumbens* (2n=4x=36) and *M. maximus* (2n=4x=32), three important species of tropical forage grasses used in pastures in tropical and subtropical areas. All these intraspecific crossings were performed by the Brazilian Agricultural Research Corporation (Embrapa) Gado de Corte, located in Campo Grande, Mato Grosso do Sul, Brazil (20°27’S, 54°37’W, 530 m), and are part of the breeding programs of this research institution.

The *U. humidicola* progeny consisted of 279 hybrids obtained from a cross between the sexual accession H031 (CIAT 26146) and the apomictic cultivar *U. humidicola* cv. BRS Tupi, as described by Vigna et al. (2016). The cross between *U. decumbens* D24/27 (sexual diploid accession tetraploidized by colchicine) and the apomict *U. decumbens* cv. Basilisk generated a progeny with 239 hybrids (Ferreira et al., 2019). Finally, the progeny of *M. maximus* included 136 hybrids originating from a cross between the sexual genotype S10 and *M. maximus* cv. Mombaça (apomictic parent) (Deo et al., 2020).

### 2.2 GBS Library Preparation

GBS libraries of the *U. decumbens* and *M. maximus* progenies were built and sequenced as described by Ferreira et al. (2019) and Deo et al. (2020), respectively. For the progeny of *U. humidicola*, DNA was extracted following Vigna et al. (2016), and GBS libraries were built according to Poland et al. (2012), containing five replicates for each parent and one for each hybrid. Genomic DNA (210 ng of DNA per individual) was digested using a combination of a rarely cutting enzyme (PstI) and a frequently cutting enzyme (MspI). Subsequently, the libraries were sequenced as 150-bp single-end reads using the High Output v2 Kit (Illumina, San Diego, CA, USA) for the NextSeq 500 platform (Illumina, San Diego, CA, USA). The quality of the resulting sequence data was evaluated using the FastQC toolkit (http://www.bioinformatics.babraham.ac.uk/projects/fastqc/).

### 2.3 GBS-SNP Discovery and Allele Dosage

We analyzed the raw data of the three biparental progenies using the Tassel-GBS pipeline (Glaubitz et al., 2014) modified for polyploids (Pereira et al., 2018b), which considers the original read depths for each SNP allele. The Bowtie2 algorithm version 2.1 (Langmead and Salzberg, 2012) was used to align the reads of the *Urochloa* spp. and *M. maximus* against the reference genomes of *Setaria viridis* v1.0 and *Panicum virgatum* v1.0, respectively, since reference genomes are not available for the species under study. In this stage, a limit of 20 dynamic programming problems (D), a maximum of four times to align a read (R) and a very-sensitive-local argument were considered. Both genomes used as references were retrieved from the Phytozome database (Goodstein et al., 2012).

For quality purposes, SNPs were submitted to a filtering procedure using VCFtools (Danecek et al., 2011), with the following parameters: maximum missing data per marker of 25% and minimum read depth of 20 reads for *M. maximus* and *U. decumbens* and 40 reads for *U. humidicola*. Due to the polyploid nature of the species, a high sequence depth is required to identify the genotypic class accurately (Cappai et al., 2020; Ferrão et al., 2020; Mollinari et al., 2020), and even higher values were used for *U. humidicola* because it is hexaploid. Finally, the Updog R package (Gerard et al., 2018) was used to estimate the allele dosage of each SNP locus, with a fixed ploidy parameter of four for *M. maximus* and *U. decumbens* and six for *U. humidicola*. The flexdog function was used with the “f1” population model for the three populations. The posterior proportion of misgenotyped individuals (prop_mis) was set at six different values (0.05, 0.1, 0.15, 0.20, 0.25 and 0.3) for *M. maximus* and *U. decumbens*, aiming to compare the rates of the tetraploid dosages in the parents and assess the influence of the number and quality of the markers in further analysis. For the hexaploid population of *U. humidicola*, prop_mis was set at 0.2.

The genotyping data were organized into marker matrices **M**_*(n x m)*_, where *n* denotes the samples, *m* denotes the markers, and the genotypes are encoded as 0, 1, 2, 3, 4, 5 or 6 for nulliplex, simplex, duplex, triplex, quadruplex, quintuplex and hexaplex data, respectively.

### 2.4 Simulated Data

Biparental F_1_ populations were simulated using the PedigreeSim R package (Voorrips and Maliepaard, 2012), a software package that simulates meiosis and uses this information to create cross populations in tetraploid species. To create the linkage map required by PedigreeSim, the previously published map for *M. maximus* (Deo et al., 2019) was used as a model to estimate the main parameters, such as the number and size of chromosomes, density, gap regions and centromere position. Eight chromosomes with sizes between 90 and 120 centimorgans (cM)and 600-900 SNP markers per chromosome, both randomly sampled, were created. In addition, the markers were considered to be distributed along the chromosomes with a minimum distance between adjacent markers of 0.1 cM. The centromere position was sampled between 10 and 50 cM, preferential pairing was set to zero, and the quadrivalent fraction was set for natural pairing. In this case, the fraction of quadrivalents arises automatically from the pairing process at the telomeres. Other options of the software were kept as default. All these files were created using R software (R Core Team, 2020).

The simulated crosses between autotetraploid parents created in PedigreeSim were based on the combinations P1 x P2, P1 x P1 (self-fertilization), P1 x P3, P1 x P4 and P3 x P4, with a progeny size of 200. The genotypes of these parents were simulated based on the rate of allele dosages of parents genotyped in real biparental progenies: P1 and P2 from *M. maximus* (Deo et al., 2020) and P3 and P4 from *U. decumbens* (Ferreira et al., 2019). Considering these rates, the haplotypes of each of the four homologous chromosomes were randomly created for each parent.

The results of the simulated crosses were converted into marker matrices (**M**), and all subsequent manipulations were performed using R software (R Core Team, 2020). To insert genotyping errors, 5% of the genotypes were randomly replaced by other genotype values with equal probability, and between 1 and 5% of the genotypes of each marker were removed to simulate missing data (NAs). Clonal individuals were simulated by duplicating the genotype of a parent, and errors and NAs were inserted as described above.

Using the tetraploid populations created in PedigreeSim software, four scenarios were established to analyze different types of contaminants that could occur in biparental populations of tropical forage grasses. The first two scenarios were represented by contaminants resulting from the reproductive mode of parents, which can reproduce by apomixis, generating clones (ACs) (1), or self-fertilization progeny of one of the parents (SPs) (2), resulting in segregating individuals. The last two scenarios represent cross-contamination, that is, when fertilization occurs by foreign pollen, resulting in half-siblings (HSs) (3), or when physical mixtures occur during seed handling, resulting in full contaminants (FCs) (4). In each of the four possible scenarios, the size of the base population was 200 hybrids (HPs), and hybrids were progressively replaced by contaminants until 25% of the samples were contaminants. In addition, to investigate a joint scenario with four parents (P1, P2, P3 and P4) with AC and SP contamination, a population of 1,200 individuals (200 P1-ACs, 200 P1-SPs, 200 HPs from P1 x P2, 200 HPs from P1 x P3, 200 HPs from P1 x P4 and 200 HPs from P3 x P4) was created. These described populations were constructed to investigate how contaminants influence PCA scatter plot dispersion patterns.

For the evaluation of the proposed contaminant identification method, 6,000 populations were simulated. Each one was composed of 200 individuals with a random number of contaminants, ranging between 1 and 50 and distributed per contaminant type considering random probabilities between 0.1 and 0.8. The populations were divided in six equal size groups according to the number of genotyped markers. Considering *n* as the total simulated markers, the groups were composed of: n/2, n/4, n/8, n/16, n/32 and n/64 markers. For each population, the subset of markers used was randomly sampled from the total simulated markers. Furthermore, a biparental population with 200 individuals (150 HP (P1 x P2), 10 AC (P1), 10 SP (P1), 10 HS1 (P1 x P3), 10 HS2 (P1 x P4) and 10 FC (P3 x P4)) was also simulated to exemplify the use of GA and CA in the contaminant identification.

### 2.5 PCA

PCAs were performed by the R package pcaMethods (Stacklies et al., 2007) using the nonlinear estimation by iterative partial least squares (NIPALS) algorithm (Wold and Krishnaiah, 1966) to calculate the eigenvalues with missing data imputation. Given a matrix X_*m,n*_ representing the n random variables (herein SNPs) across m individuals, this analysis transforms X by multiplying it by the orthogonal eigenvectors, generating a matrix X_*m,p*_ of new p variables (the PCs) with specific mathematical properties (Maćkiewicz and Ratajczak, 1993). The ggplot2 R package (Wickham and Chang, 2016) was used to construct scatter plots of the first two PCs. These graphical visualizations were used to identify clustering patterns that may be associated with contaminants in the progeny.

### 2.6 GA

The term GA is employed here to refer to an analysis that evaluates all samples of a progeny considering what is genotypically expected for a contaminant. Three different measures were created for evaluating the samples: GA-I for AC identification and GA-II for SP identification, both accounting for a similarity rate between the sample and one of the parents (computed separately for each), and GA-III, accounting for a rate of unexpected genotypes in each sample considering the genotypes of both parents, enabling the identification of HSs and FCs in the progeny.

To investigate whether an individual *x* is an AC of a parent *p*, the GA-I scores were calculated using the marker matrix **M** with *n* rows (individuals) and *m* columns (markers). Then, the similarity between *x* and *p* was the proportion of allele dosages in M_*x,i*_ that satisfied the condition M_*x,i*_ = M_*p,i*_ with 1 ≤ *i* ≥ *m*. This measure is based on the presumption that, given Mendel’s law, each individual inherits genetic material from its parents (Mendel, 1866; Miko, 2008). However, if one of the parents reproduces through apomixis, a genetically identical progeny is produced (Hand and Koltunow, 2014). Therefore, in a suite of Mendelian loci, if a putative individual shows a high similarity (GA-I close to 1.00) with one of the parents, it can be considered a clone of this parent.

In the case of SP samples, the GA-II scores were calculated by computing the similarity between the progeny samples and the parents considering only nulliplex allele dosages; i.e., for a parent p and an individual x, GA-II was the proportion of allele dosages in M_*x,i*_ (with 1 ≤ *i* ≤ *m*) that satisfied M_*x,i*_ = M_*p,i*_ = 0. If a parent reproduces through self-fertilization, Mendelian segregation should be observed. Using a tetraploid species as an example, a parent with the genotype AABB at a specific locus, after self-fertilization, would generate a progeny with genotypes in all possible doses (AAAA, AAAB, AABB, ABBB and BBBB) (Hackett et al., 2013). However, if we focus only on markers for which the parent had a nulliplex genotype (AAAA), the progeny produced would be genetically identical to the parent at those loci. Thus, GA-II computes a similarity rate between the sample and the parent considering only those markers; in this situation, it was expected that SP contaminants would present GA-II scores close to 1.00.

For the HSs and FCs, the GA-III term calculates the rate of unexpected allele dosages for the progeny individuals across all the markers. Considering the combination of gametes for parent *p1* and *p2* at a SNP *i* with 1 ≤ *i* ≥ *m*, the GA-III of an individual *x* is the proportion of unexpected allele dosages for its set of markers. Considering the allele dosage of each parent at each marker, it is possible to define which dosage is not expected in their progeny. For example, if one parent is nulliplex (AAAA) for a marker and the other is simplex (AAAB), the gametes produced by the nulliplex are all AA, and for the simplex, they can be AA or AB (Hackett et al., 2013). Their combination can produce a progeny only nulliplex or simplex for this marker, and the presence of other dosage types is evidence that this individual may not belong to the cross. In this way, for all markers, GA-III tested whether the sample’s genotype was expected considering both parental genotypes, computing a rate of unexpected genotypes for each sample. In this analysis, it was expected that HSs and FCs would show higher GA-III scores than HPs, enabling their identification.

### 2.7 CA

The contaminant identification process is based on a CA performed using an average linkage hierarchical clustering approach with R software (R Core Team, 2020). Considering the GAs calculated, pairwise Euclidean distances between these values were calculated across the progeny and used to obtain 27 different clustering indexes (Supplementary Table 1) with numbers of clusters varying from 2 to 15, implemented in the R package NbClust (Charrad et al., 2014). The package automatically calculates the indexes, defines the best clustering scheme and classifies the samples into clusters.

Contaminant identification was then performed with the best clustering configuration scheme. Individuals in groups separated from most of the population were considered contaminants and classified according to the following rules applied to the GA measures within these clusters: (1) individuals within a cluster having the greatest GA-I values for one parent were considered ACs when the median of these measures was greater than 0.75; (2) individuals within a cluster with the median GA-II values for one parent greater than 0.75 and not belonging to group (1) were considered SPs; and (3) individuals not belonging to groups (1) and (2) and with a within-group minimum GA-III value greater than the maximum measure of the group with the minimum GA-III median were considered HSs/FCs. Therefore, in a simplified and automated process with no ad hoc decisions, we obtained the final data set with parents and their corresponding true hybrids.

The method was evaluated on a set of 1,000 simulated populations, computing the following metrics considering the most indicated clustering scheme: mean rate of hybrids correctly identified (MRH), mean rate of contaminants correctly identified (MRC), mean rate of ACs correctly identified (MRAC), mean rate of SPs correctly identified (MRSP) and mean rate of cross-contaminants (HSs/FCs) correctly identified (MRCC). Furthermore, the same metrics were computed considering the three most indicated clustering schemes; in this situation, the highest rate among the three schemes for each simulated population was used to calculate the mean.

### 2.8 Contaminant Identification in Real Data

Combining GA, CA and PCA, we established a four-step contaminant identification methodology as follows, and applied it to the real populations (Figure 1, Part III Contaminant Identification):

**Figure 1.**
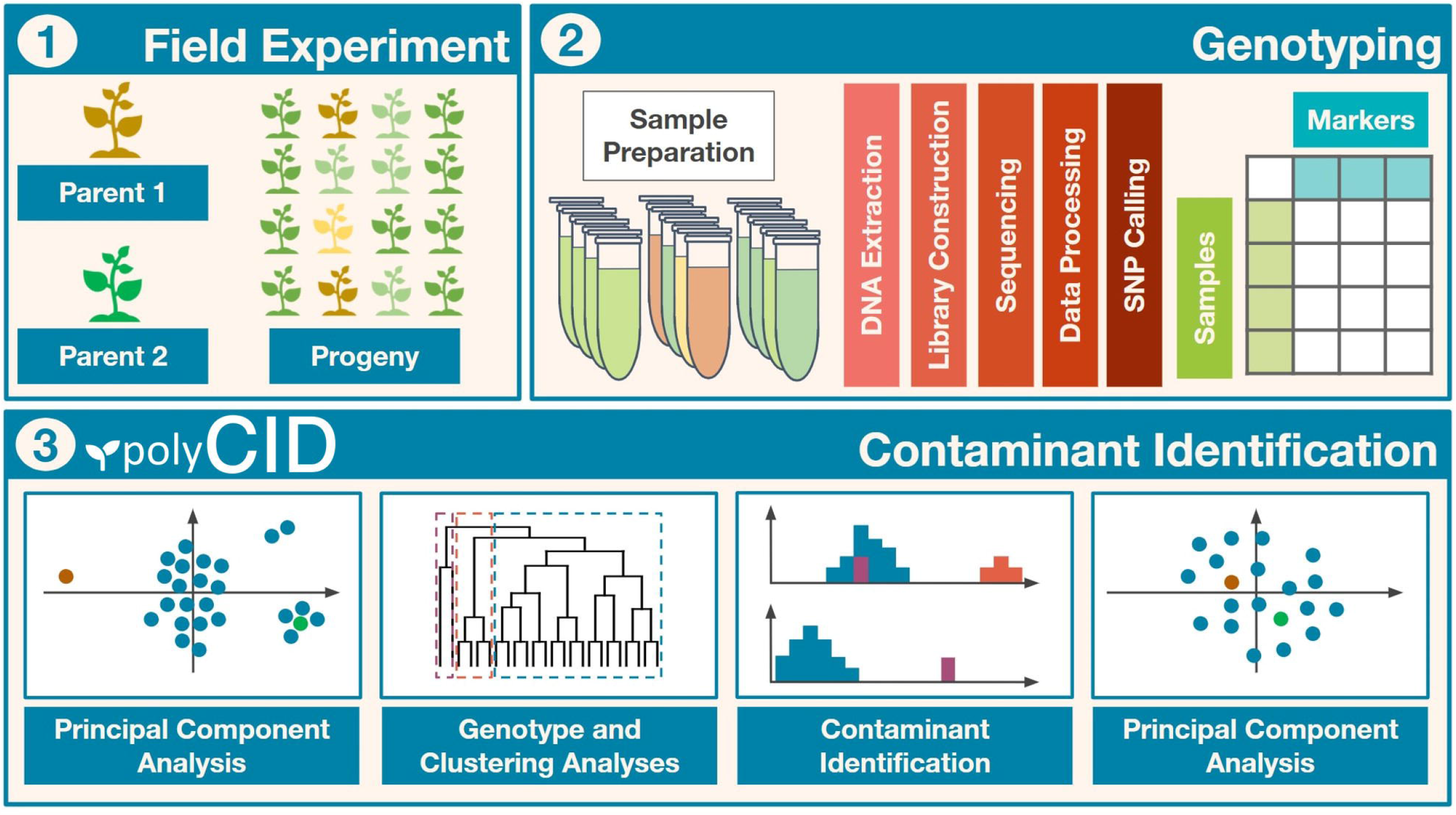
Workflow of the contaminant identification process. (I) Field experiment to obtain the biparental progeny. (II) Population genotyping and bioinformatic analyses. (III) Methodology with principal component analysis (PCA), genotype analysis (GA) and clustering analysis (CA) to identify and remove contaminants.

1. Construction of a scatter plot with the first PCs from a PCA performed with the SNP data organized according to allele dosage, looking for evidence of contaminants in the population. When no contaminants are detected, simulated clones (25% of the population) from one of the parents can be artificially added to the population, changing the dispersion pattern of individuals and inducing contaminant separation;
2. Calculation of five different GA measures for each individual (GA-I and GA-II, considering parents 1 and 2, respectively, and GA-III). GAI and GAII were calculated in the same way for all ploidy, but for hexaploid progeny, GAIII was adapted considering its respective segregation;
3. Performance of CA to identify the putative contaminants, classified according to the GA value differences described in Section 2.7 (AC, SP, HS or FC);
4. Removal of the identified contaminants from the population, and performance of a new PCA and construction of a biplot to confirm the expected dispersion pattern of a population with no contaminants.

All these procedures were unified in polyCID Shiny app, created using R software together with the libraries shiny (Chang et al., 2021), shinydashboard (https://cran.r-project.org/web/packages/shinydashboard/index.html) and DT (https://cran.r-project.org/web/packages/DT/index.html). polyCID is an R-Shiny Web graphical user interface (GUI) that combines all the described analyses in a simple way and provides a user-friendly tool, fully available and documented at https://github.com/lagmunicamp/polycid.

## 3 Results

The results are organized as follows. First, the genotyping and allele dosage information for the three biparental progenies of the tropical forage species are presented (3.1). Next, the application of PCA to the simulated data is discussed (3.2). Then, the use of GA and CA in contaminant identification in the simulated data is described (3.3), and finally, the results obtained from contaminant identification in real data are presented (3.4).

### 3.1 GBS-SNP Discovery and Allele Dosage Estimation

After SNP calling using the Tassel-GBS pipeline (Glaubitz et al., 2014) modified for polyploids (Pereira et al., 2018b), filtering markers for missing data (NAs) and read depth with VCFtools (Danecek et al., 2011), we obtained 15,279 SNP markers for *U. humidicola*, 8,036 for *U. decumbens* and 6,337 for *M. maximus*. Three individuals (“Bh181”, “Bh226”, and “Bh245”) of the *U. humidicola* progeny were removed because of the high content of missing data (> 44%).

The Updog R package (Gerard et al., 2018) was used to estimate the allele dosage for the SNP loci identified in each progeny. For the six values of prop_mis used (0.05, 0.10, 0.15, 0.20, 0.25 and 0.30), 4,003, 5,179, 5,863, 6,068, 6,161 and 6,215 markers were obtained for *M. maximus* and 1,195, 1,745, 2,303, 2,862, 3,165 and 3,243 markers were obtained for *U. decumbens*, respectively, while 7,253 markers were obtained for *U. humidicola* using a prop_mis value of 0.20.

### 3.2 PCA

Marker matrices of each simulated scenario were used to perform a PCA, looking for patterns spanned by the first two PCs that can aid in the identification of contaminant samples. Details of these simulated scenarios, such as the size of chromosomes, position of centromeres and number of markers, can be found in the supplementary material (Supplementary Tables 2-4). The PCA scatter plot of the simulated population without contaminants had hybrids and parents distributed with no apparent clustering patterns among the individuals, with 4% of variance explained by the first two PCs (Supplementary Figure 1).

The same biplot distribution was observed when only one contaminant was added to the biparental population, i.e., an AC (Figure 2A), SP (Supplementary Figure 2), HS (Supplementary Figure 3) or FC (Supplementary Figure 4). In these situations, the genetic variation related to contamination could not be detected by the first components and therefore assessed by visual inspection. When the number of contaminants was progressively increased in the scenarios, the dispersion pattern of the scatter plots began to reveal separation of the contaminants from the hybrids. For the scenarios, five (Figure 2B), four (Supplementary Figure 5), six (Supplementary Figure 6) and three (Supplementary Figure 7) contaminants were necessary to clearly visualize the separation. Adding these contaminants changed the source of variation in the first PCs, which changed little (<0.2%). As the number of contaminants increased to 25% of the population, it was possible to observe in the PCA biplot that the hybrids were projected between the parents, the ACs/SPs were closer to the parent of origin (Figure 2C and Supplementary Figure 8), the HSs/FCs formed separated groups (Supplementary Figures 9 and 10), and the sums of variance in the first two PCs changed to values between 10.8% and 17%.

**Figure 2.**
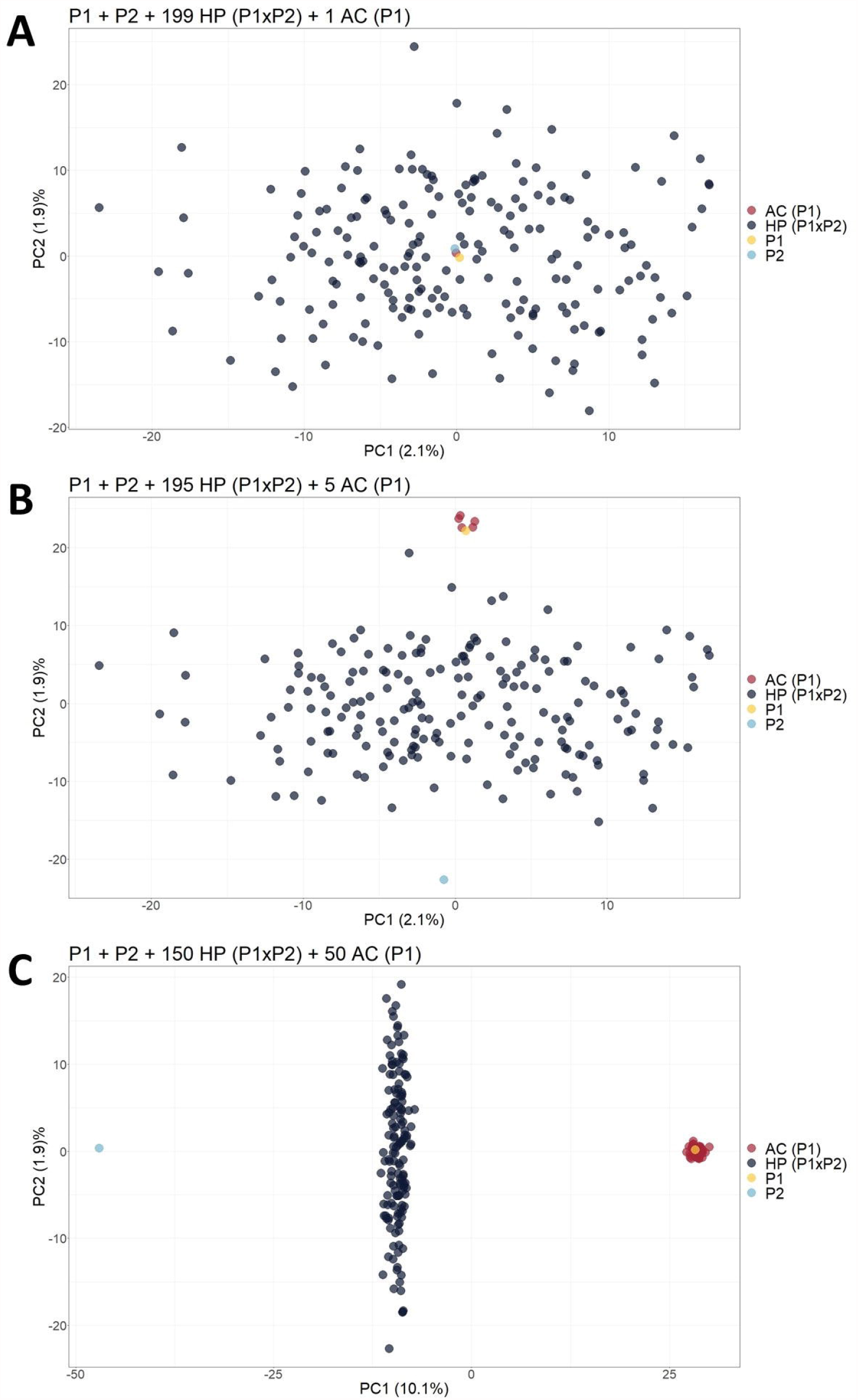
Principal component analysis (PCA)-based scatter plots showing the change in dispersion pattern as the apomictic clone (AC) of P1 increases in frequency in the simulated population. (**A**) Progeny with 199 hybrids (HPs) and 1 AC; (**B**) Progeny with 195 HPs and 5 ACs; (**C**) Progeny with 150 HPs and 50 ACs. The axis represents the first and second principal components, explaining 2.1% and 1.9% of the variance, respectively, for (**A**), 2.1% and 1.9% for (**B**) and 10.1% and 1.9% for (**C**).

Considering that the analysis of the first two PCs through a PCA scatter plot could not reveal contaminants at low frequencies, biparental populations with 199 HPs and one contaminant were simulated, and 50 ACs (25%) of one of the parents were included. This unique contaminant was an SP (Scenario 2), AC (Scenario 3) or FC (Scenario 4). As a result, we observed that the inclusion of these simulated clones, which occurs in real populations, changed the sums of variance in the first two PCs to a value of approximately 10.3% and increased the dispersion pattern in the PCA scatter plot, leading to the formation of different subgroups that allowed the visualization of SP or HS contaminants (Supplementary Figures 11 and 12). On the other hand, FCs and HPs were grouped together and could not be identified visually in the scatter plot (Supplementary Figure 13).

Finally, when simulating an open pollination population with four different possible parents, the biplot of the PCs was able to provide visual separation of the different progenies. It was possible to identify each cross since HPs formed a subgroup between their respective parents. In addition, the AC and SP contaminants grouped together with their parents (Figure 3).

**Figure 3.**
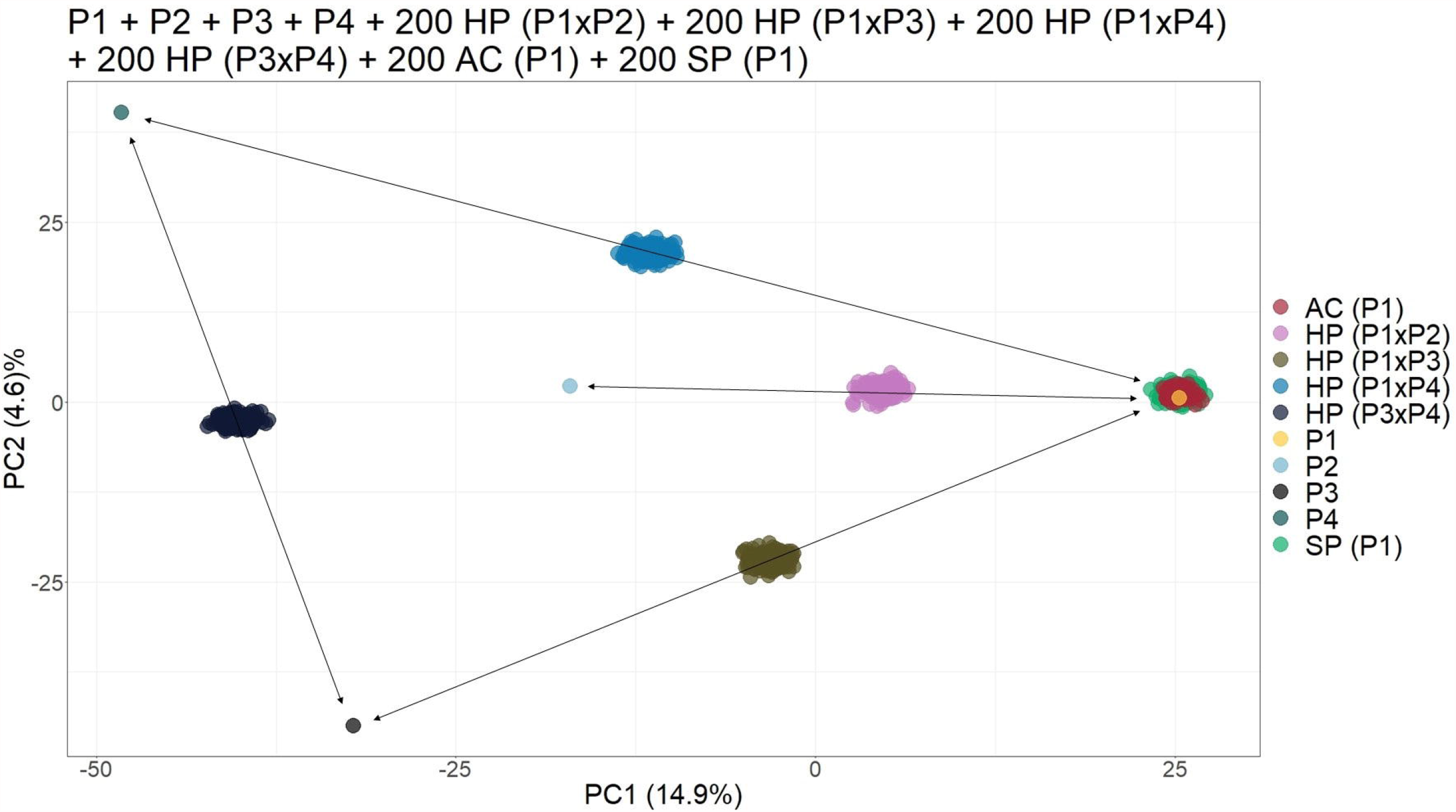
Principal component analysis (PCA)-based scatter plot showing the simulated population composed of four parents (P1, P2, P3 and P4), 200 apomictic clones - ACs (P1), 200 self-fertilization progeny - SPs (P1), 200 half-siblings - HSs (P1 x P2), and 200 hybrids - HPs (P1 x P3, P1 x P4 and P3 × 43). The axis represents the first and second principal components, with 14.9% and 4.6% of the variance explained, respectively.

### 3.3 Automatic Contaminant Identification

To look for patterns in contaminant GA measures, the three described GAs were calculated in a simulated population of 200 samples composed of 150 HP and 50 contaminants (10 ACs, 10 SPs, 10 HS1s, 10 HS2s and 10 FCs); thus, five different values for each putative hybrid were generated. We analyzed how GA histograms behave for each type of contamination. In Figure 4A, AC individuals formed a group with the greatest GA-I scores for Parent 1 (red circle) and were removed to analyze the other histograms. In the same way, the GA-II histogram (Figure 4B) showed that the SP samples had the highest scores for Parent 1 (red circle), and these individuals were also removed. Finally, in Figure 4C, the GA-III histogram showed that the HP samples had lower scores than the HS and FC contaminants. For correct hybrid definition, these contaminants were also removed to generate a proper hybrid data set.

**Figure 4.**
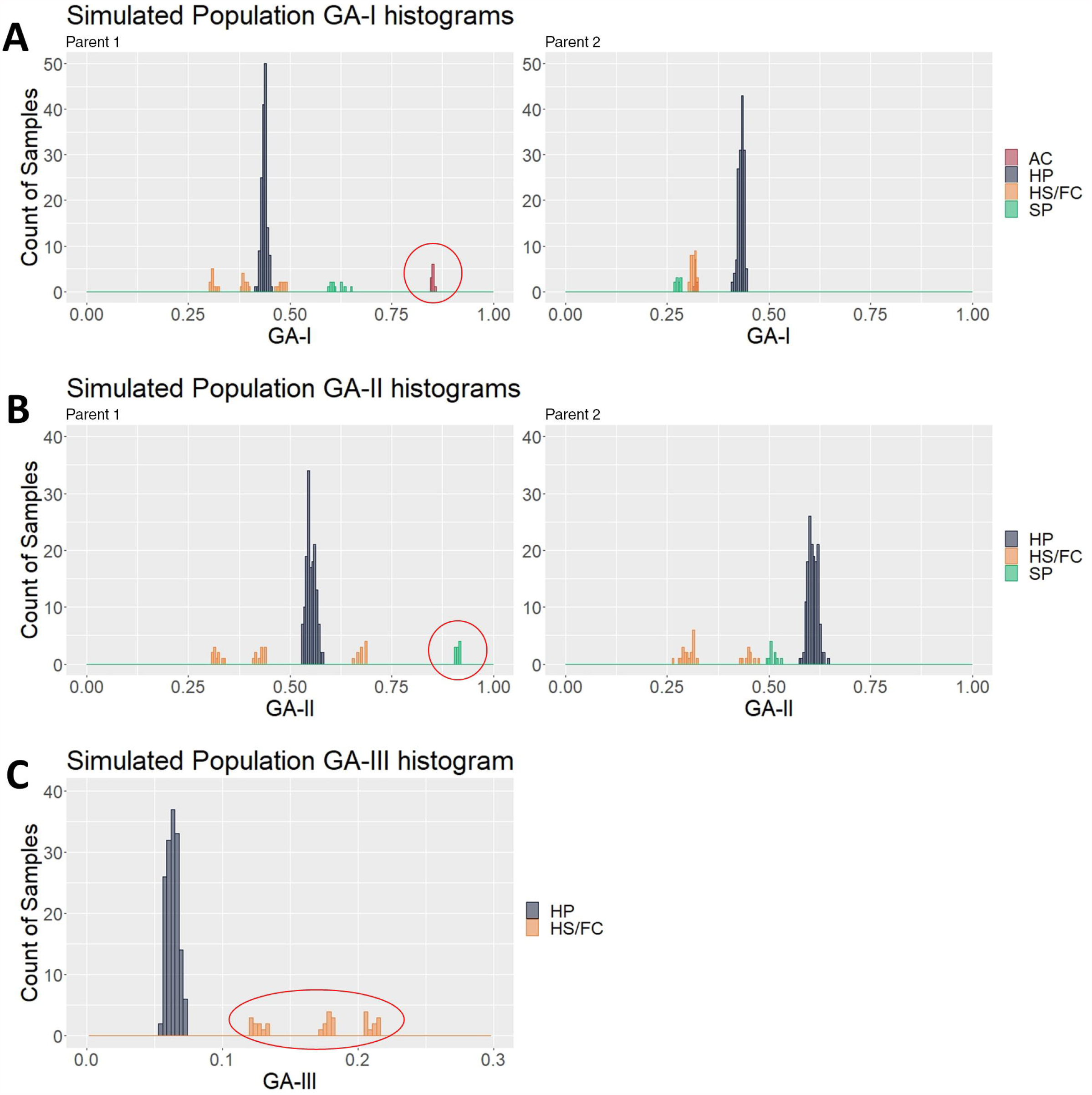
Genotype analysis (GA)-based histograms of the simulated population with samples classified in the clustering analyses (CA). (**A**), (**B**) and (**C**) show the results from GA-I, GA-II and GA-III, respectively. Red circles highlight the identified contaminants, i.e., apomictic clones (AC), self-progeny (SP) and half siblings/full contaminants (HS/FC), in contrast to the hybrid progeny (HP).

By using the idea underlying these visual histogram inspections, we implemented on GA measures a clustering-based approach for an automatic contaminant identification. The established methodology employs a single hierarchical clustering algorithm on a different range of cluster numbers, defining the best clustering scheme with 27 clustering indexes (Supplementary Table S1). Employing this approach on the simulated population previously described, we observed that the defined CA separated the samples in six different groups: one for the HP and five for each type of contaminant, exactly corresponding to the simulated categories (Figure 4). Therefore, we evaluated its accuracy on additional 6,000 simulated populations, and checked its appropriateness using six set sizes of markers in a broad range of possible contamination scenarios. The sets were of the following sizes: 2,758, 1379, 689, 344, 172 and 86 markers. For each marker’s set size, 1,000 populations were simulated sampling markers from a total of 5,516.

The mean rate of hybrids correctly identified (MRH) was 100% for all sets of markers, except for the smallest one (86 markers), which had a slight reduction. On the other hand, the MRC was around 90% for the three largest sets (2,758, 1,379 and 689 markers), but then started decreasing, reaching the value of 48% in the smallest one (Figure 5-A). It was possible to observe that the methodology failed only for the smallest set (86 markers), in which a true hybrid was considered a contaminant, but it rarely discarded reliable data. Regarding the contaminant classification and considering the largest sets of markers, 69%, 72% and 84% were observed for MRAC, MRSP and MRCC, respectively. Then, we observed a slight reduction in the 689 markers set, which showed values of 63% (MRAC), 66% (MRSP) and 76% (MRCC), followed by 49% (MRAC), 50% (MRSP) and 15% (MRCC) in the smallest set (Figure 5-A).

**Figure 5.**
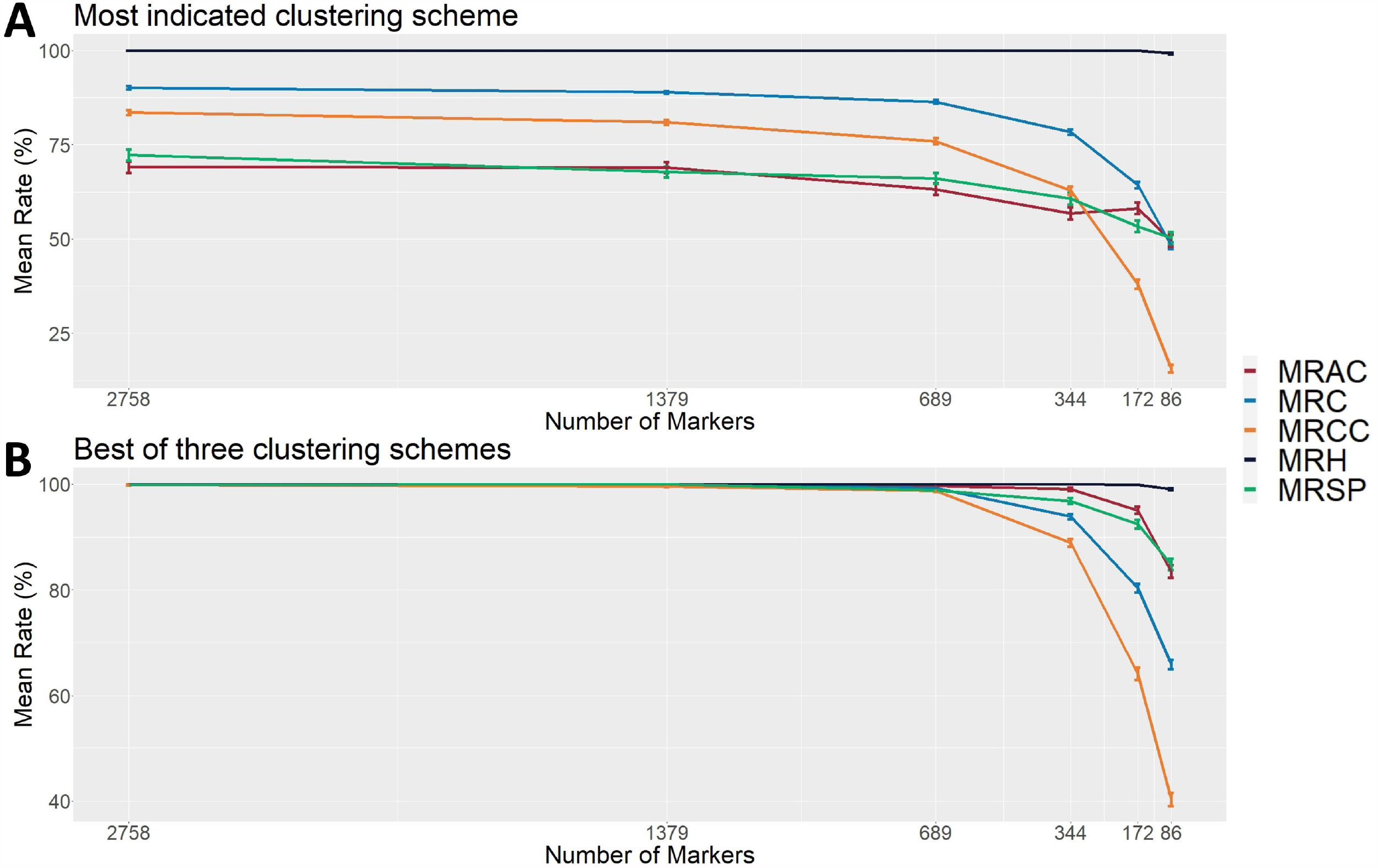
Evaluation metrics of the proposed methodology considering six different set sizes of makers in simulated populations (1,000 populations for each set). (A) the most indicated clustering scheme, and (B) the best out of the first three most indicated schemes found with the 27 employed clustering indexes. The indicated rates are the mean rate of hybrids correctly identified (MRH), the mean rate of contaminants correctly identified (MRC), the mean rate of apomictic clones correctly identified (MRAC), the mean rate of selfed-progeny correctly identified (MRSP), and the mean rate of cross-contaminants (half-sibling/full contaminant) correctly identified (MRCC).

In function of these modest values, we also evaluated the method efficiency on the second and third best clustering configurations identified by the calculated indexes. Considering the best group separation within these three possible configurations also noticed in GA histograms, we achieved a performance improvement in all set markers, reaching an approximate accuracy of 100% in the three largest sets for all types of samples. Next, we observed a slight reduction (to values higher than 90%) in the set of 344 markers, and more prominent reductions in the two smallest sets, reaching the values of 49% (MRAC), 85% (MRSP) and 40% (MRCC) (Figure 5-B). These findings suggest that, in real applications, such evaluations in these three cluster configurations may represent an additional step for increasing the method reliability.

### 3.4 Contaminant Identification in Real Populations

After investigations with simulated populations, the proposed methodology was applied to real genotyping data from three biparental F1 progenies of tropical forage grasses. For the progeny of *M. maximus*, the PCA plots with different values of prop_mis showed similar sample dispersion patterns and a reduction in variance explained by the first two PCs from 10.1% to 7.5% as the number of markers increased. Therefore, the dataset obtained with the default value of prop_mis = 0.20 was used in further analysis. Even though there was no clear group formation in the PCA scatter plot, the pattern of parents on opposite sides and HPs grouped between them provided evidence of the presence of contaminants (Supplementary Figure 14A). Similarly, in the PCA with simulated ACs, these two individuals remained close to Parent 2 (*M. maximus* cv. Mombaça) (Supplementary Figure 14B)

The CA with GA revealed two clusters in the *M. maximus* progeny, with 134 and two samples. The GA-I histogram for Parent 2 (*M. maximus* cv. Mombaça) showed that the cluster with two samples had high scores and must be considered putative ACs of Parent 2 (*M. maximus* cv. Mombaça) (Supplementary Figure 15A). On the other hand, GA II and III showed no evidence of other types of contaminants in the *M. maximus* progeny (Supplementary Figure 15B and C). The exclusion of these two individuals resulted in a PCA scatter plot with the expected pattern (Supplementary Figure 14C).

For the progeny of *U. decumbens*, the PCA biplots for the different values of prop_mis showed different sample dispersion patterns (data not shown). As this is a very intuitive measure for the quality of SNPs when estimating allele dosage (Gerard et al., 2018), we chose to be conservative and use the most restrictive filter, 0.05, ensuring the selection of markers with high quality. The first PCA scatter plot showed strong evidence of contaminants in the population (Figure 6A). The algorithm found three clusters with 184, 49 and three samples. In the GA histograms, both clusters 2 and 3 had high GA-I scores for Parent 2 (*U. decumbens* cv. Basilisk), providing evidence that those samples were putative ACs of this parent (Figure 7A). The other GA histograms showed no evidence of other contaminants (Figure 7B and C). Once again, after the elimination of these ACs, the PCA scatter plot showed the expected pattern for progeny without contaminants (Figure 6B).

**Figure 6.**
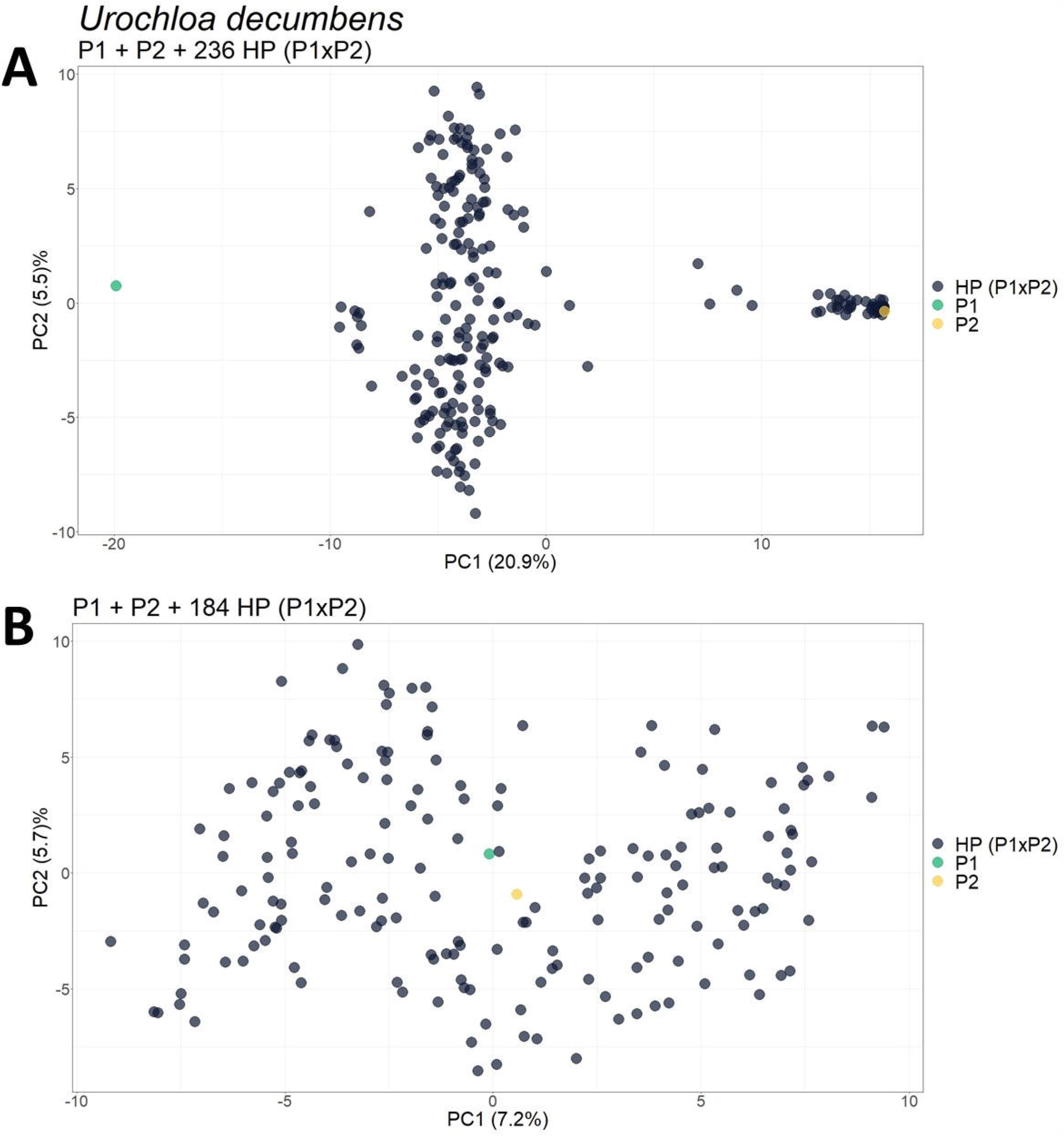
Principal component analysis (PCA)-scatter plots showing the *U. decumbens* progeny for the set of SNP markers filtered by 0.05 for *prop_mis*. (**A**) Original population composed of two parents (P1 and P2) and their progeny of 236 hybrids (HP (P1xP2)); (**B**) Population without the 52 apomictic clones (AC) identified. The axis represents the first and second principal components, explaining 20.9% and 5.5%, respectively, of the variance for (**A**) and 7.2% and 5.7% for (**C**).

**Figure 7.**
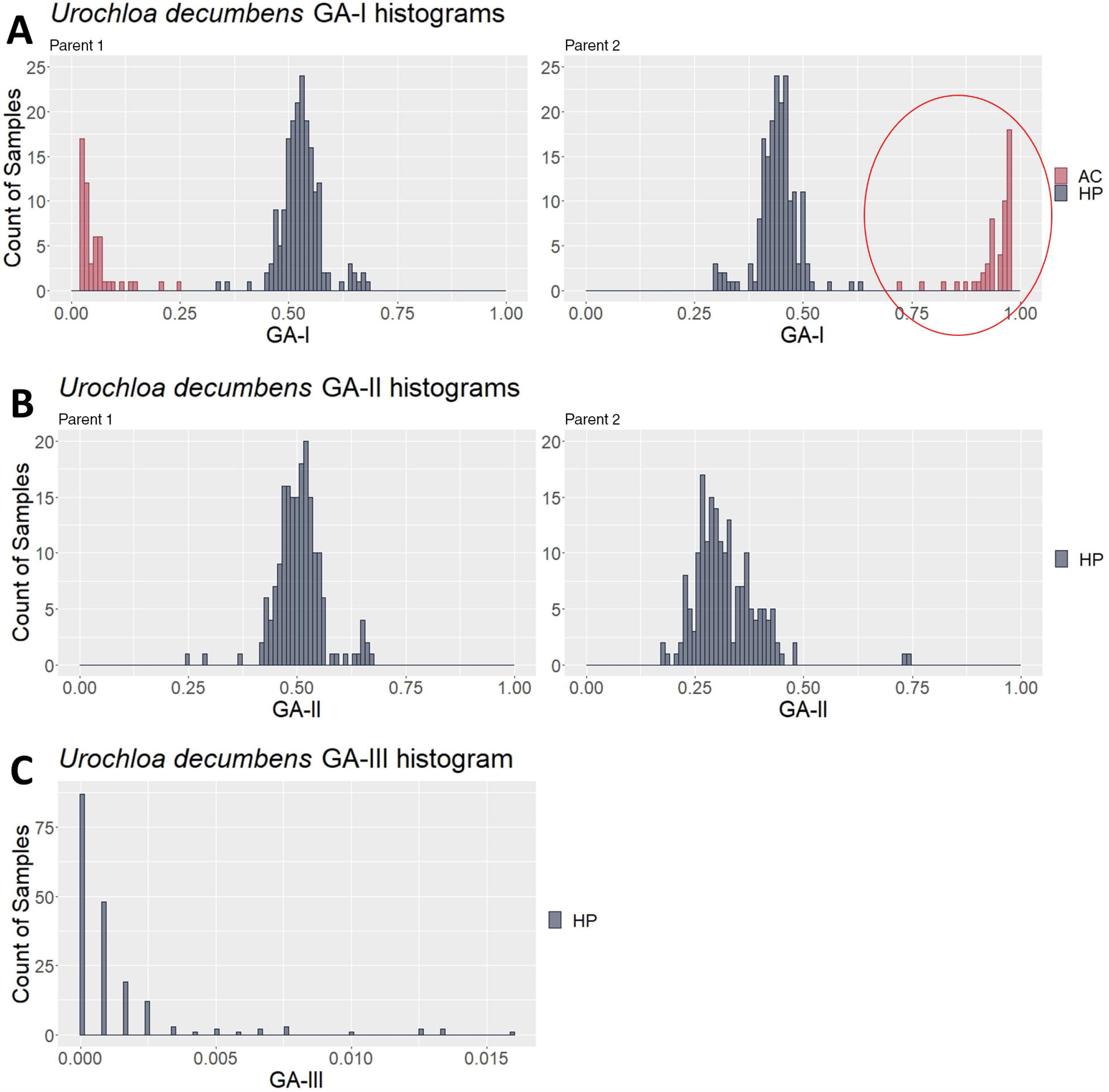
Genotype analysis (GA)-based histograms of *Urochloa decumbens* with samples classified in the clustering analysis (CA). (**A**), (**B**) and (**C**) show the results from GA-I, GA-II and GA-III, respectively. The red circle highlights the identified apomictic clone (AC) contaminants in contrast to the hybrid progeny (HP).

For the hexaploid biparental population of *U. humidicola*, the scatter plot of the first PCs showed strong evidence of the presence of AC and/or SP contaminants (Supplementary Figure 16A). The clustering analysis of the GA scores separated the progeny into two clusters with 211 and 65 samples. The histogram of GA-I for Parent 1 (*U. humidicola* H031) showed that the cluster with 65 samples had scores close to 1.0 (Supplementary Figure 17A), representing putative ACs of the respective parent. GA-II and GA-III showed no evidence of contaminants (Supplementary Figure 17B and C). Finally, the PCA without the previously identified contaminants also showed the expected pattern for progeny without contaminants (Supplementary Figure 16B).

### 3.5 polyCID Shiny App

Finally, we implemented the polyCID Shiny app, a Web GUI that provides all previously described analyses in a user-friendly tool that allows users to identify contaminants in biparental progeny in a simple way. polyCID is completely R based and easy to install and presents a graphical interface designed for nonexpert users, with several functions for interactive visualization of the results. The package accepts SNP data in the form of marker matrices with allele dosage information, loads this information and performs the four-step contaminant identification methodology, as described in Section 2.8. The Shiny-based GUI is included in the package as a standalone application, available at https://github.com/lagmunicamp/polycid.

## 4 Discussion

Experimental populations used in breeding programs are usually derived from a controlled cross between two or more parents, but depending on the field experiment, the species analyzed and its reproductive biology, individuals may be generated from mixtures of seeds, foreign pollination during open pollinated crosses, self-fertilization or apomixis by one of the parents during the crosses. The nonidentification of these contaminant individuals can interfere not only in the selection cycles of breeding programs (Telfer et al., 2015) but also in studies of genetic diversity (Ji et al., 2013), population structure (Alam et al., 2018), linkage mapping (Deo et al., 2020) and association mapping (Laucou et al., 2018) since it can generate biased results.

In most available studies, the identification of contaminants involved the use of microsatellites and morphological markers, but this strategy can be costly and time consuming (Santos et al., 2014; Jha et al., 2016; Zhao et al., 2017; Patella et al., 2019), especially for polyploid species. In these cases, progeny evaluation is often performed using few microsatellite markers in polyacrylamide gels, and frequently, other analyses are needed, such as genetic distance analysis (Santos et al., 2014). The low number of microsatellite markers, usually in the tens or hundreds, may prevent the identification of contaminants. Considering this scenario, we used GBS (Poland et al., 2012) to identify thousands of SNP markers with allele dosage information and to propose a methodology that facilitates the identification of contaminants in biparental crossbreeding of polyploid species. Despite the emergence of several pipelines for the analysis of GBS data in polyploids, the application of these markers in parentage analysis is still little explored.

Currently, several software packages can deal with genetic data to assign paternity or parentage to individuals at the diploid level using microsatellite or SNP markers and likelihood-based or Bayesian methods (Kalinowski et al., 2007; Jones and Wang, 2010; Anderson, 2012; Huisman, 2017), in addition to other approaches (Hayes, 2011; Heaton et al., 2014; Grashei et al., 2018; Whalen et al., 2019). For polyploids, the few resources available are limited to microsatellite data (Spielmann et al., 2015; Zwart et al., 2016). Another common approach in polyploids is to estimate pairwise relatedness (r) (Huang et al., 2015; Amadeu et al., 2020), for example, to assess the relationships between parents and offspring and full-sibs and half-sibs in progenies. In addition, identity-by-descent (IBD) has been used to assess the probabilities of inheritance of particular combinations of parental haplotypes (Zheng et al., 2016), which are also quite difficult to evaluate in polyploid progenies. For both approaches, the parameters are estimated in a pairwise manner, and the results are evaluated for each pair, making the analysis even more laborious.

For breeding programs that make use of biparental crosses, the major challenge is to precisely identify whether there are contaminating individuals to be excluded from the progeny (Martuscello et al., 2009; Ma and Amos, 2012; Santos et al., 2014; Subashini et al., 2014; Simeão et al., 2016b; Matias et al., 2019; Deo et al., 2020). In this context, no studies have proposed a unified pipeline focused on identifying the most common contaminants in biparental crossings, especially in polyploid species, and supplying such a pipeline is the main objective of this work. Therefore, we propose an unprecedented automatized pipeline that is based on PCA, GA and CA to identify and classify all types of contaminants in a biparental progeny. The proposed methodology was developed and tested in F_1_ biparental crosses of tropical forage grasses, but it can be applied to any tetraploid or hexaploid species since the parents of the F_1_ biparental cross are known.

### 4.1 Contaminant Identification in Simulated Data based on PCA, GA and CA

PCA is a multivariate data technique used to represent a dataset as orthogonal variables named PCs. Aiming at reducing the dimensionality of a set of variables through linear combinations, repeated information can be removed while the maximum variance-covariance structure of these variables is maintained (Jolliffe and Cadima, 2016). As the first two components explain the most variance in the SNP data, a scatter plot of the samples in a Cartesian plane with these PCs is a way to visually identify similarities and differences and determine whether samples can be grouped (Ringnér, 2008). Our results showed that in a simulated biparental F_1_ progeny without the presence of contaminants, the first components showed a two-dimensional pattern in which the population was distributed between the two parents (Supplementary Figure 1), which was expected since these individuals were closely related. As the first PCs generally reflect the variance related to the population structure in the sample, individuals from the same population form a unique cluster in a subspace spanned by the first two eigenvectors (Ma and Amos, 2012).

Considering the four simulated scenarios described above, a contaminant frequency of approximately 3% in a progeny is needed to observe a different pattern of PCs that allows the identification of contaminants (Figure 2B and Supplementary Figures 5-7), which shows the inefficiency of employing a PCA biplot for such an approach. In cases with a lower percentage of ACs, SPs or HSs, duplicating the genotype of one of the parents to generate artificial clones proved to be an alternative way to change the dispersion pattern of individuals, inducing the projection of contaminants as separated from the real hybrids (Supplementary Figures 11 and 12). This occurred because the values for the linear combination increased for the PC1 vector, and the source of variation changed to be based on the presence of inserted clones. On the other hand, we found that FC contamination could be detected with fewer contaminating individuals (1.5% contaminants in relation to the total population) due to the different genetic backgrounds in relation to the progeny. This high genetic variability modifies the first components and thereby facilitates the identification of FCs in the PCA.

In general, PCA has a better-defined pattern that allows more inferences about population relationships, not at the individual level (Patterson et al., 2006). It has been widely performed using microsatellite and SNP markers for diploid and polyploid species to evaluate population structure (Larsen et al., 2018; Lara et al., 2019), to infer genetic ancestry (Byun et al., 2017), to predict genomic breeding values (Macciotta et al., 2010), and for other applications. However, for contaminant identification, the use of the first components from PCA, even those successfully employed in forage grass polyploids (Lara et al., 2019; Deo et al., 2020), proved to be insufficient in most scenarios; therefore, other approaches are required.

In the pipeline described here, we propose the use of PCA as a first step to visualize the data and suggest the presence of possible contaminants in biparental crosses, evaluated through appropriate CAs. The main limitation of PCA lies in cases with few contaminants, i.e., less than 3% of the progeny, which has already been reported in tropical forage grasses (Deo et al., 2020). Artificially inserting simulated clones from one of the parents changed the dispersion pattern in most cases; however, when the contaminants were HSs or FCs, the variance was still not captured by the first components. Therefore, PCA itself could not effectively identify and classify the contaminants and, for this reason, was combined with other techniques. We suggested the use of specific GA measures as inputs for CAs as a methodological workflow capable of accomplishing this task.

The fundamental idea underlying GA-I, GA-II and GA-III was to identify incompatibilities between putative hybrids and their parents as a strategy to conclusively demonstrate their parentage. For such analyses, it is expected that the GA scores from different populations (here, hybrids and contaminants) form different distributions with specific parameters. Although there are other approaches for parentage estimation already discussed in the literature, such as Identity by Descent (IBD) or pairwise relatedness (r) (Huang et al., 2015; Zheng et al., 2016; Amadeu et al., 2020), these measures indicate how close an individual is to another in a given population, regardless of the degree of relationship. GA measures, on the other hand, differ from these in terms of their focus on the genetic relationships in biparental populations for which both parents are known. Here, the main objective is to compute scores that are related to the type of contaminants expected in such populations, enabling not only identification but also classification.

In all simulated populations with 689 or more genotyped markers, the proposed methodology could correctly identify and classify almost 100% of the samples, ratifying the appropriateness of the proposed pipeline. The size of the markers set employed in different scenarios has been demonstrated to have a large effect on the accuracy of the methodology, as we observed a positive correlation between the two variables. Nevertheless, considering the most indicated clustering scheme, sets with more than 689 markers did not cause an expressive accuracy increase (Figure 5). Previous studies have evaluated accuracies in function of number of markers in different genomic approaches, such as parentage assignment and genomic selection, and found similar results (Wang et al, 2012; Arruda et al, 2015; Lenz et al, 2017; Whalen et al, 2018). However, finding and generalizing the optimal number of markers for this methodology is complicated because it may be influenced by various factors, including the species, population size, contaminants quantity/type and sequencing/genotyping techniques.

Even though the CA identifies different groups of individuals with similar GA measures, the association of each group with a contaminant type requires an additional step, which is important because identifying the type of contamination (in the case of AC or SP contamination) can assist the breeder in better understanding the reproductive biology of the species or genotype. On the other hand, identifying HS or FC contaminants highlights the need for greater control during the field experiment, avoiding foreign pollen or seed mixtures. Interestingly, we noticed that each cluster captured a distinct pattern in the GA measures, a phenomenon that can be leveraged to decipher the contaminant origin of the individuals. Importantly, by using the proposed approach, we did not find any configuration in which true hybrids were discarded, which is of great value for real applications.

In summary, our proposal is a unique methodology that brings together all types of contamination in a single identification pipeline, representing an important resource for breeders, who need specific tools to deal with such contamination. Instead of relying solely on the putative population structure revealed by PCA methodologies, GA indexes are calculated, taking into account the genetics behind the origin of the contaminants. Compared to the exclusive use of PCA, this pipeline identifies one or a few contaminating individuals with more confidence. This increased confidence makes this methodology ideal for situations in the field that lead to mixtures of seeds or foreign pollen during fertilization, which usually occur at low rates.

### 4.2 Contamination Identification in Real Data

PCA, GA and CA using genotypic data from the *M. maximus, U. decumbens* and *U. humidicola* F_1_ progenies led to the conclusion that these real progenies had AC contaminants (Supplementary Figures 16 and 19 and Figure 7). For *M. maximus*, the two detected clones (1.4% of the population) corroborated the findings of Deo et al. (2019), while for *U. decumbens*, 52 individuals (21.7% of the population) were identified as clones of the male parent. It is possible that these clones were inserted into these two progenies during seed collection. Additionally, the male parent was used as a control in the field experiments, and the plants may have produced seeds and/or seedlings that became mixed with the real progeny. As the female parent of these populations were entirely sexual, the absence of SPs suggests the predominance of allogamy in these plants and self-incompatibility as the main mechanism to guarantee this mode of reproduction.

We extended this methodology for the identification of contaminants in hexaploid species, represented in this study by *U. humidicola* (2n=6x=36). GA-I and GA-II were performed in the same way as for tetraploid species, but GA-III was adapted considering the segregation and possible combination of gametes in hexaploid species. For the progeny of *U. humidicola*, the combined PC and GA-I histogram analysis suggested the presence of 61 clones of the female parent (21.8% of the population). This result suggests that the genotype H031 (CIAT 26146) also reproduces through facultative apomixis, even though it has been widely cited in the literature as a unique obligate sexual genotype of *U. humidicola* (Jungmann et al., 2010; Vigna et al., 2016). It is known that the expression of apomixis in the same genotype may vary with the flowering season in other grasses (Rios et al., 2013). It might be that the mode of reproduction of H031 was evaluated at the end of flowering or under a specific environmental condition, when the proportion of sexuality was greater than apomixis; therefore, this genotype might be a facultative apomict with high rates of sexuality (Karunarathne et al., 2020). In addition, the sexual genotypes of the *Urochloa* spp. can present a certain degree of SI (Keller-Grein et al., 1996; Dusi et al., 2010), and Worthington et al. (2019) reported the detection of 12 individuals derived from accidental self-pollination of *U. humidicola* H031. Therefore, there is a need to enrich the current understanding of *U. humidicola* biology and reproduction mode, which are important for developing suitable breeding and selection methods (Barcaccia and Albertini, 2013).

All three forage progenies used in this work have already been used in studies previously developed for the construction of genetic maps. Deo et al. (2020) identified and removed two contaminants in *M. maximus* progeny by PCA, which were also identified as contaminants by our methodology. However, for the progeny of *U. decumbens* (Ferreira et al., 2019) and *U. humidicola* (Vigna et al., 2016), only an analysis of the bands of the hybrids identified by genotyping with dozens of microsatellites and random amplified polymorphic DNA (RAPD) markers (Bitencourt et al., 2008), respectively, was performed, and no clones could be identified through this approach. Therefore, the absence of an adequate methodology and/or sufficient number of markers for the prior identification of contaminants has resulted in genetic maps constructed with genetic information including some false hybrids, and consequently, these maps may contain bias that should be considered by researchers.

Our methodology proved to be useful in practical situations of breeding programs of tropical forage grasses, including the identification of different progenies from multiparent crosses, which may be extended to other polyploid crops. The identification of contaminants in early stages of breeding cycles can greatly increase the efficiency of programs, preventing costs with false hybrids that might otherwise only be discarded in later phases of selection. Conversely, it allows the size of the useful population to increase, optimizing the breeding populations. Although the use of molecular markers is not yet a reality in many breeding programs, it is important to assess potential expenses brought by false hybrids, which might surpass the cost of large-scale genotyping technologies (such as GBS), which have been experiencing considerable cheapening in recent years. PCA, GA and CA were combined in a simple and automated pipeline, and the coupling of a low-cost genotyping with such pipeline thus allows a more precise and efficient detection of incompatibilities between a group of putative hybrids and the identification of contaminants in biparental crosses of tetraploid and hexaploid species.

The implementation of this simple approach in the identification of contaminants in biparental progenies of polyploid species can increase the efficiency of breeding programs. In this context, the polyCID Shiny app was designed to enhance the ability of breeders to use our methodology, even with no bioinformatics expertise. Great advances in sequencing technologies and genotyping tools have enabled us to explore vast amounts of genetic data in a more cost effective and faster way; however, the ability to handle and apply this genome information to breeding remains a significant barrier for most breeders and experimental researchers. Therefore, we designed the polyCID Shiny app as an interactive and user-friendly application that is completely R based and easy to install, incorporating the analysis in a single environment and enabling users to extract information on contaminant individuals without requiring knowledge of a programming language.

Finally, although our analyses were performed with real and simulated progenies of tropical forage grasses, this methodology can be extended to any biparental progeny of tetraploid or hexaploid species. It can be applied in the early stages of genomic studies with GBS in biparental polyploid progenies, such as genetic linkage map construction and genomic prediction, to identify possible contaminants. However, as the price of SNP genotyping is constantly decreasing and other polyploid genotyping tools are emerging, the application of our methodology even in experiments that do not involve SNPs may be possible, mainly in intermediate and final stages of the breeding program to confirm the absence of contamination in the final stages and cultivar release. In the case of genotyping with a lower number of molecular markers, it is suggested that simulation studies be carried out a priori, taking into account how the number and quality of the markers affect the final results.

## Author Contributions

FBM, ACLM, AHA, BBZV and APS conceived the study. LC, RMS, SCLB, MFS, LJ and CBV conducted the field experiments. ACLM, RCUF and BBZV performed the laboratory experiments. FBM, ACLM and AHA analyzed the data. FBM, ACLM, AHA, RCUF and BBZV wrote the manuscript. AHA and FBM implemented the Shiny web app. All authors read and approved the manuscript.

## Acknowledgments

We would like to acknowledge the Fundação de Amparo à Pesquisa de do Estado de São Paulo (FAPESP), the Conselho Nacional de Desenvolvimento Científico e Tecnológico (CNPq), and the Coordenação de Aperfeiçoamento de Pessoal de Nível Superior (CAPES). We also acknowledge the Brazilian Agricultural Research Corporation (Embrapa Gado de Corte) for providing the populations used in this study. This manuscript was previously posted to bioRxiv at https://www.biorxiv.org/

## Data Availability Statement

The raw sequence data for *U. humidicola* have been submitted to the NCBI Sequence Read Archive (SRA) under accession number PRJNA703438. Raw GBS data for *U. decumbens* and *M. maximus* were previously submitted to the NCBI database under accession numbers SRP148665 and PRJNA563938, respectively.

