## Supplementary Material for "An Automated SNP-Based Approach for Contaminant Identification in Biparental Polyploid Populations of Tropical Forage Grasses"

#### 1 Supplementary Methods

##### 1.1 Clustering Indices

**Supplementary Table 1.** Overview of the indices implemented in the NbClust package.

| **Name of the index in NbClust** | **Name of the index in the literature** | **References** |
|---|---|---|
| "kl" | KL | (Krzanowski and Lai, 1988) |
| "ch" | Calinski and Harabasz | (Calinski and Harabasz, 1974) |
| "hartigan" | Hartigan | (Hartigan, 1975) |
| "ccc" | Cubic clustering criterion (CCC) | (Sarle, 1983) |
| "scott" | n log (\|T\|/\|W\|) | (Scott, 1971) |
| "marriot" | $k^2$ \|W\| | (Marriot, 1971) |
| "trcovw" | Trace Cov W | (Milligan and Cooper, 1985) |
| "tracew" | Trace W | (Edwards and Cavalli-Sforza, 1965; Friedman and Rubin, 1967) |
| "friedman" | Trace $W^{-1}B$ | (Friedman and Rubin, 1967) |
| "rubin" | \|T\|/\|W\| | (Friedman and Rubin, 1967) |
| "cindex" | C-index | (Hubert and Levin, 1976) |
| "db" | Davies and Bouldin | (Davies and Bouldin, 1979) |
| "silhouette" | Silhouette | (Rousseeuw, 1987) |
| "duda" | Je(2)/Je(1) | (Duda and Hart, 1973) |
| "pseudot2" | $\text{Pseudot}^2$ | (Duda and Hart, 1973) |
| "beale" | Beale | (Beale, 1969) |
| "ratkowsky" | $c/k^{.5}$ | (Ratkowsky and Lance, 1978) |
| "ball" | Ball and Hall | (Ball and Hall, 1965) |
| "ptbiserial" | Point-Biserial | (Kraemer, 1982) |
| "gap" | Gap | (Tibshirani et al., 2001) |

| | | |
|---|---|---|
| "mcclain" | McClain and Rao | (Mcclain et al., 1975) |
| "gamma" | Gamma | (Baker and Hubert, 1975) |
| "gplus" | G(+) | (Rohlf, 1974) |
| "tau" | Tau | (Rohlf, 1974) |
| "dunn" | Dunn | (Dunn, 1974) |
| "sdindex" | SD | (Halkidi et al., 2000) |
| "sdbw" | SDbw | (Halkidi et al., 2001) |

### 2 Supplementary Results

#### 2.1 Simulation Details

The constructed linkage map used in PedigreeSim software consists of 8 chromosomes with 5,516 randomly distributed SNPs; the length, centromere position and number of markers of each chromosome can be found in Supplementary Table 1. Furthermore, as the software requires the parental genotypes for the markers to simulate the crosses, the *M. maximus* and *U. decumbens* parental allele dosage rates were used to genotype the simulated parents (P1, P2, P3 and P4), and the actual number of markers for each dosage in the set of 5,516 markers is shown in Supplementary Tables 2 and 3.

**Supplementary Table 2.** Information on the linkage map created for the simulation analysis.

| Chromosome | Size (cM) | Position of the centromere (cM) | Number of markers |
|---|---|---|---|
| **1** | 108 | 44 | 665 |
| **2** | 98 | 50 | 615 |
| **3** | 101 | 17 | 643 |
| **4** | 99 | 43 | 644 |
| **5** | 115 | 27 | 685 |
| **6** | 110 | 45 | 766 |
| **7** | 112 | 39 | 611 |
| **8** | 95 | 29 | 887 |

**Supplementary Table 3.** Rates of each allele dosage used to genotype the simulated parents. P1 and P2 were based on the parents of the *M. maximus* progeny, and P3 and P4 were based on the parents of the *U. decumbens* progeny.

| | Nulliplex (0) | Simplex (1) | Duplex (2) | Triplex (3) | Quadruplex (4) |
|---|---|---|---|---|---|
| **P1** | 0.4253 | 0.3899 | 0.1694 | 0.0110 | 0.0044 |
| **P2** | 0.3061 | 0.5255 | 0.1440 | 0.0213 | 0.0031 |
| **P3** | 0.5430 | 0.3008 | 0.1429 | 0.0098 | 0.0035 |
| **P4** | 0.1559 | 0.4930 | 0.2802 | 0.0671 | 0.0038 |

**Supplementary Table 4.** Number of SNP markers classified in each of the five possible dosage classes for the simulated parents, totaling 5,516 markers.

| | Nulliplex (0) | Simplex (1) | Duplex (2) | Triplex (3) | Quadruplex (4) |
|---|---|---|---|---|---|
| **P1** | 2,447 | 2,090 | 897 | 56 | 26 |
| **P2** | 1,652 | 2,876 | 833 | 138 | 17 |
| **P3** | 3,002 | 1,686 | 769 | 42 | 17 |
| **P4** | 848 | 2,705 | 1,552 | 388 | 23 |

#### 2.2 Principal Component Analysis of Simulated Populations

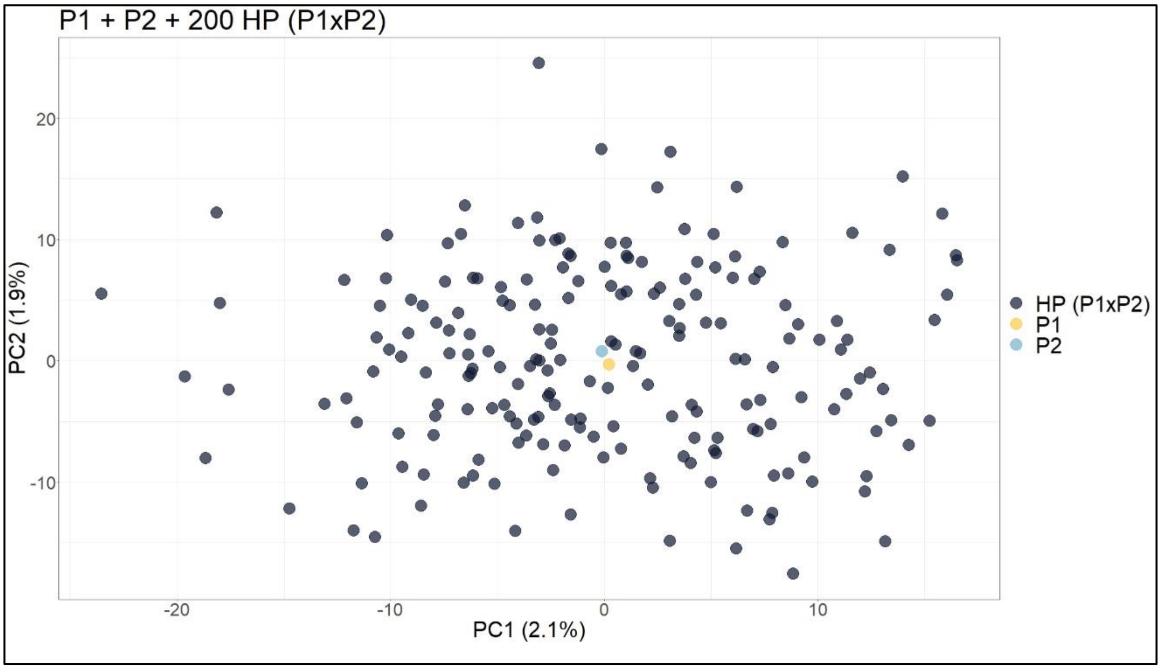


**Supplementary Figure 1**. Principal component analysis scatter plot showing the simulated population with two parents (P1 and P2) and 200 hybrids (HP (P1xP2)). The axes represent the first and second principal components, which explain 2.1% and 1.9% of the variance, respectively.

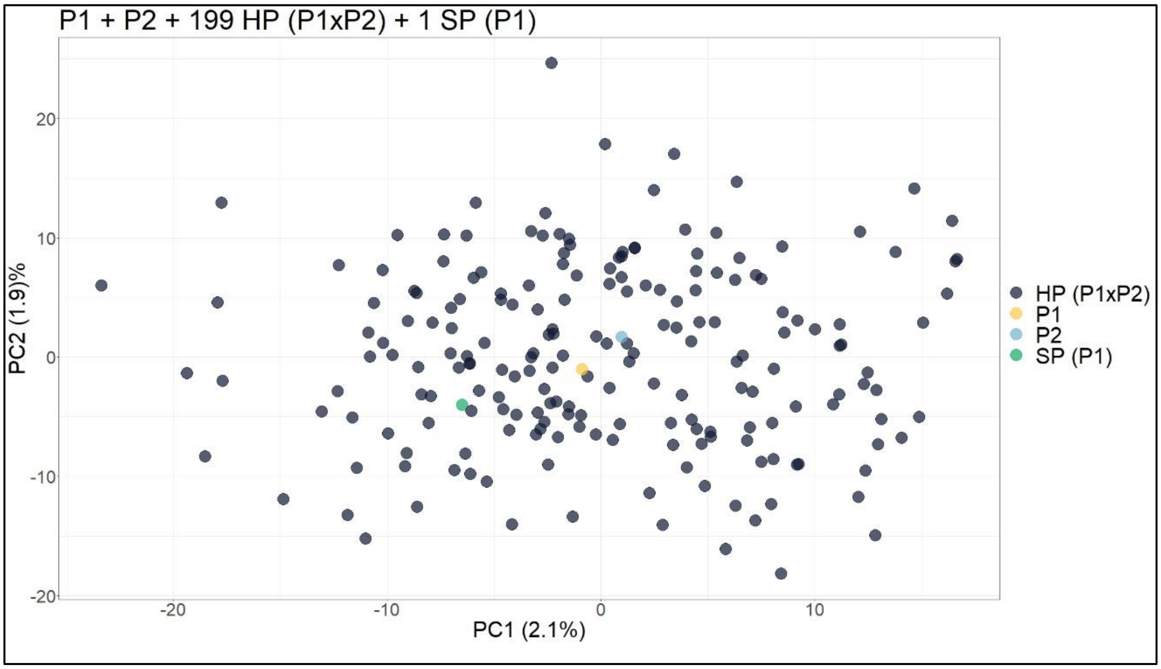


**Supplementary Figure 2**. Principal component analysis scatter plot showing the simulated population with two parents (P1 and P2), 199 hybrids (HP P1xP2) and one self-fertilization progeny (SP (P1xP1)). The axes represent the first and second principal components, which explain 2.1% and 1.9% of the variance, respectively.

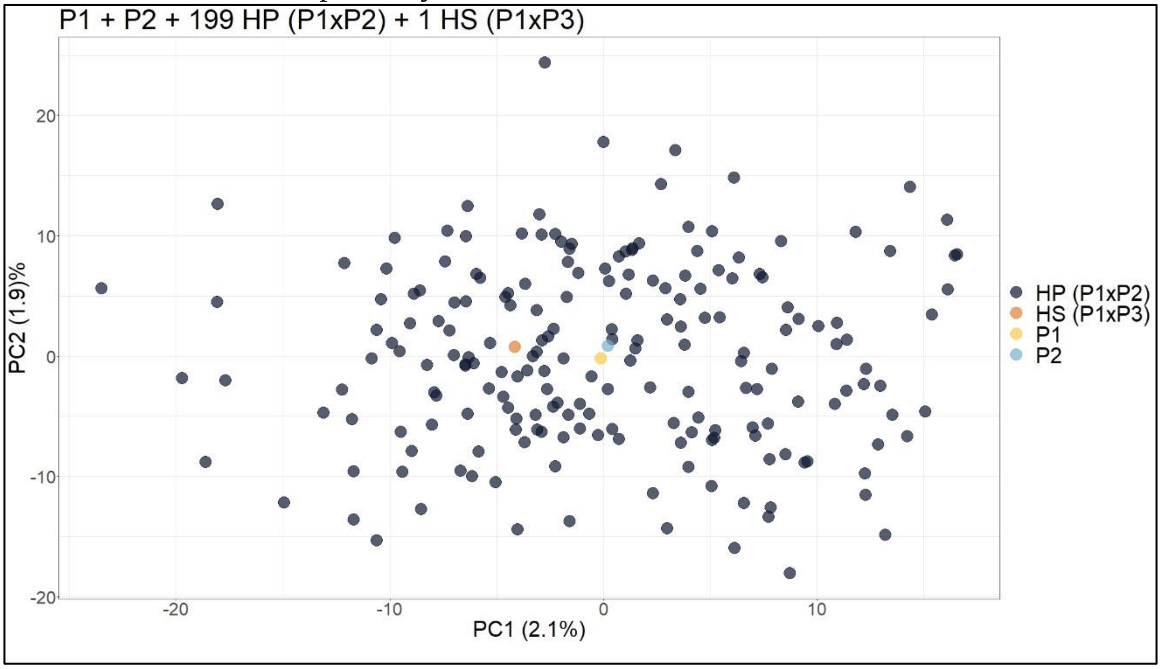


**Supplementary Figure 3**. Principal component analysis scatter plot showing the simulated population with two parents (P1 and P2), 199 hybrids (HP (P1 x P2)) and one half-sibling (HS (P1 x P3)). The axes represent the first and second principal components, which explain 2.1% and 1.9% of the variance, respectively.

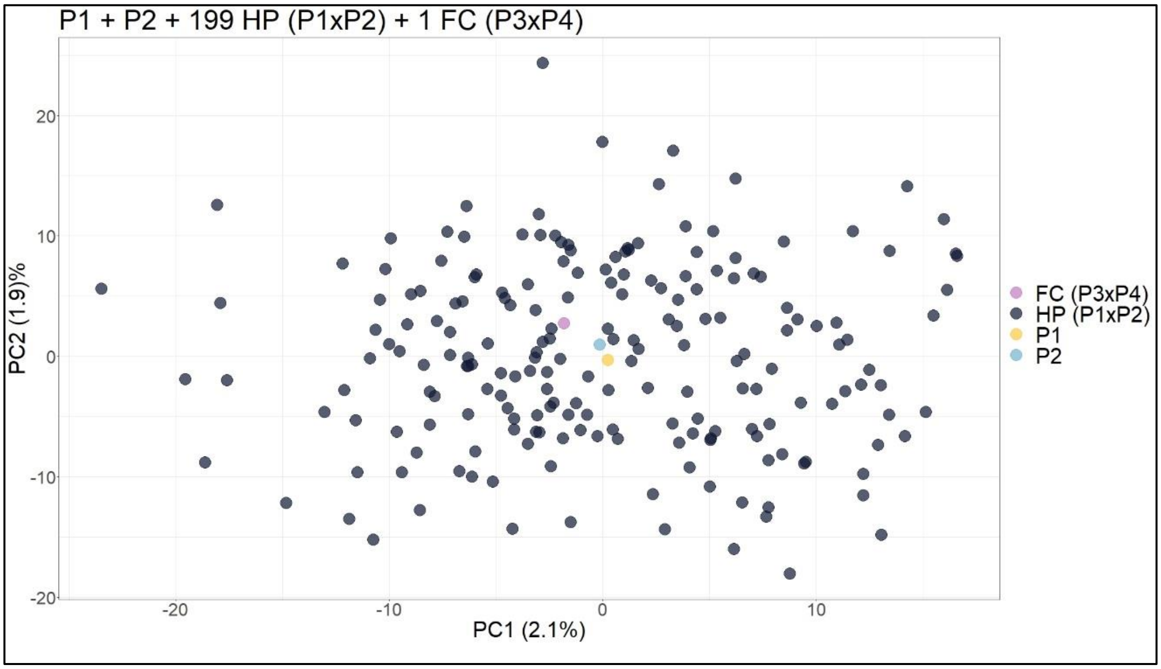


**Supplementary Figure 4**. Principal component analysis scatter plot showing the simulated population with two parents (P1 and P2), 199 hybrids (HP (P1 x P2)) and one full contaminant (FC (P3 x P4)). The axes represent the first and second principal components, which explain 2.1% and 1.9% of the variance, respectively.

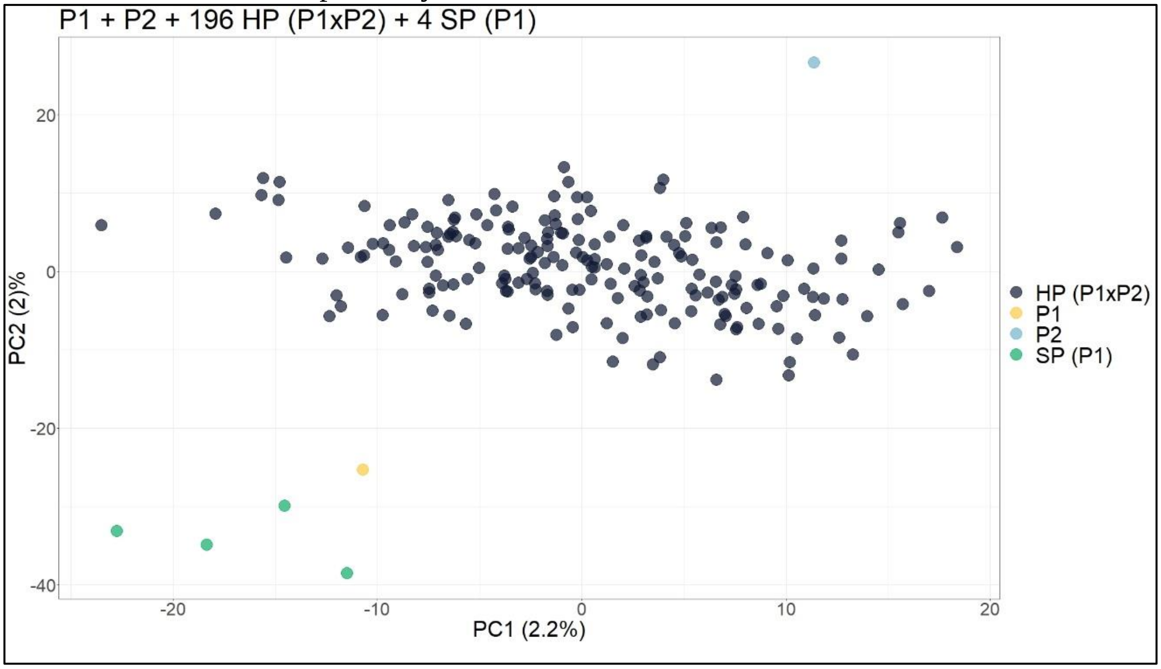


**Supplementary Figure 5**. Principal component analysis scatter plot showing the simulated population with two parents (P1 and P2), 196 hybrids (HP (P1 x P2)) and four self-fertilization progenies (SP (P1 x P1)). The axes represent the first and second principal components, which explain 2.2% and 2% of the variance, respectively.

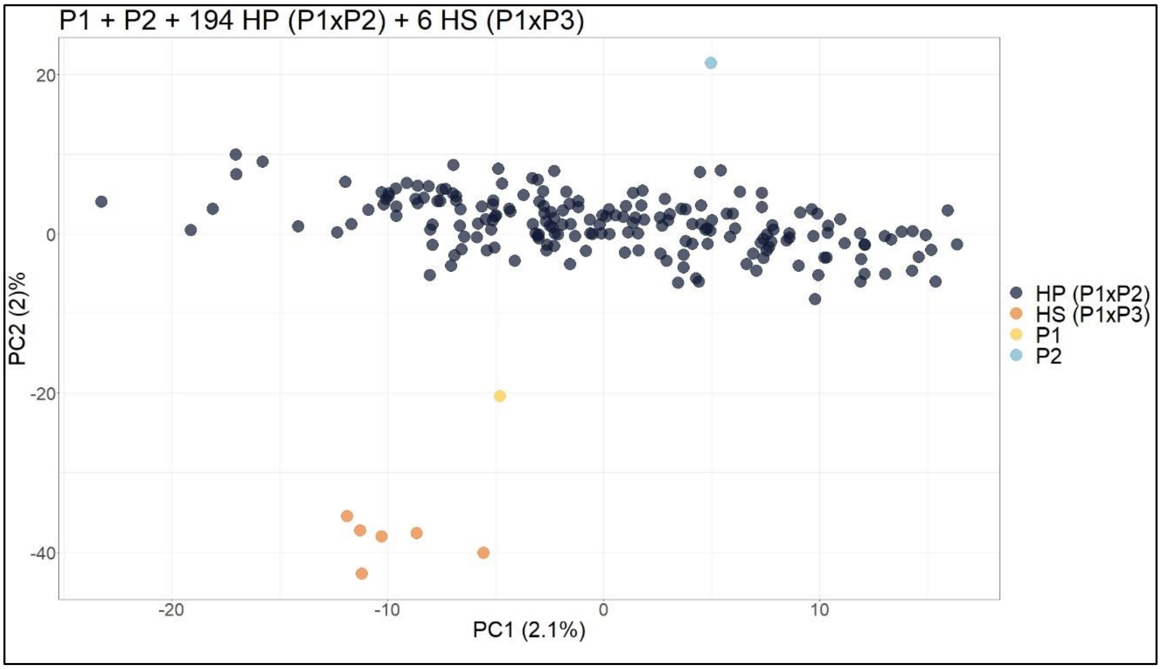


**Supplementary Figure 6**. Principal component analysis scatter plot showing the simulated population with two parents (P1 and P2), 194 hybrids (HP (P1 x P2)) and six half-siblings (HS (P1 x P3)). The axes represent the first and second principal components, which explain 2.1% and 2% of the variance, respectively.

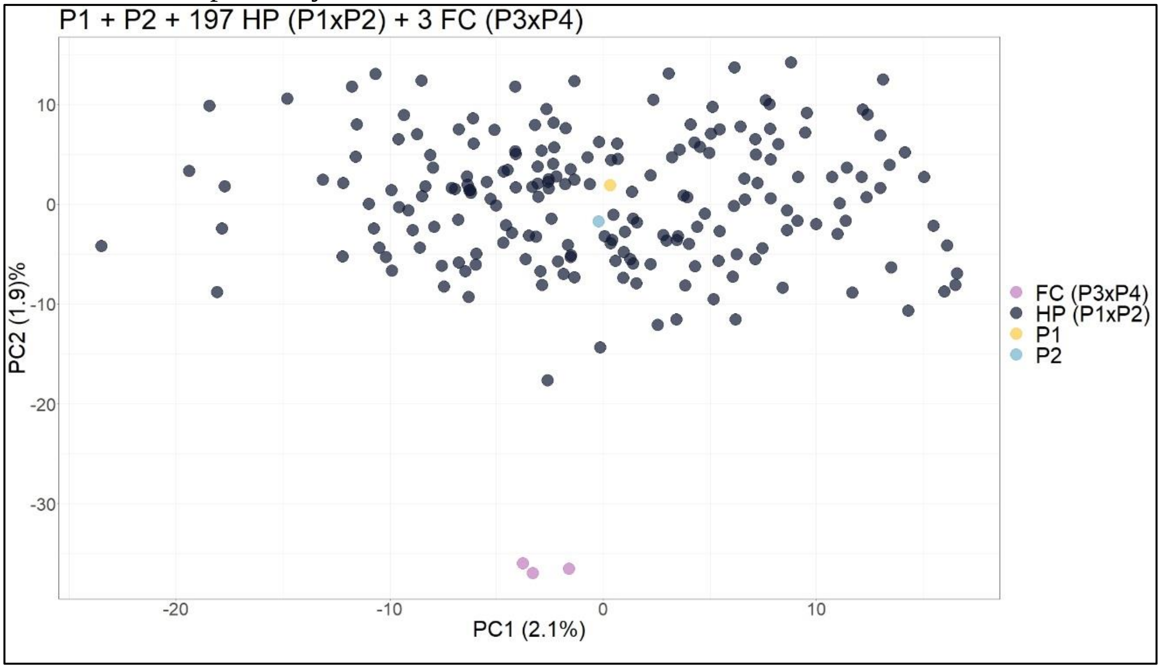


**Supplementary Figure 7**. Principal component analysis scatter plot showing the simulated population with two parents (P1 and P2), 197 hybrids (HP (P1 x P2)) and three full contaminants (FC (P3 x P4)). The axes represent the first and second principal components, which explain 2.1% and 1.9% of the variance, respectively.

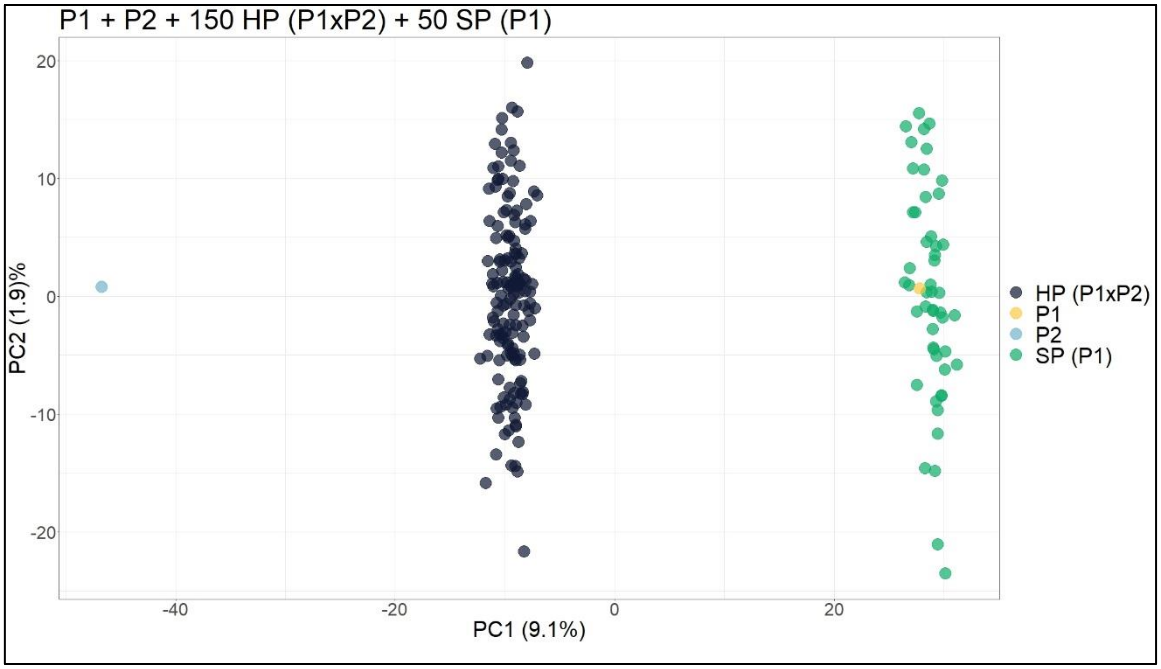


**Supplementary Figure 8**. Principal component analysis scatter plot showing the simulated population with two parents (P1 and P2), 150 hybrids (HP (P1 x P2)) and 50 self-fertilization progenies (SP (P1 x P1)). The axes represent the first and second principal components, which explain 9.1% and 1.9% of the variance, respectively.

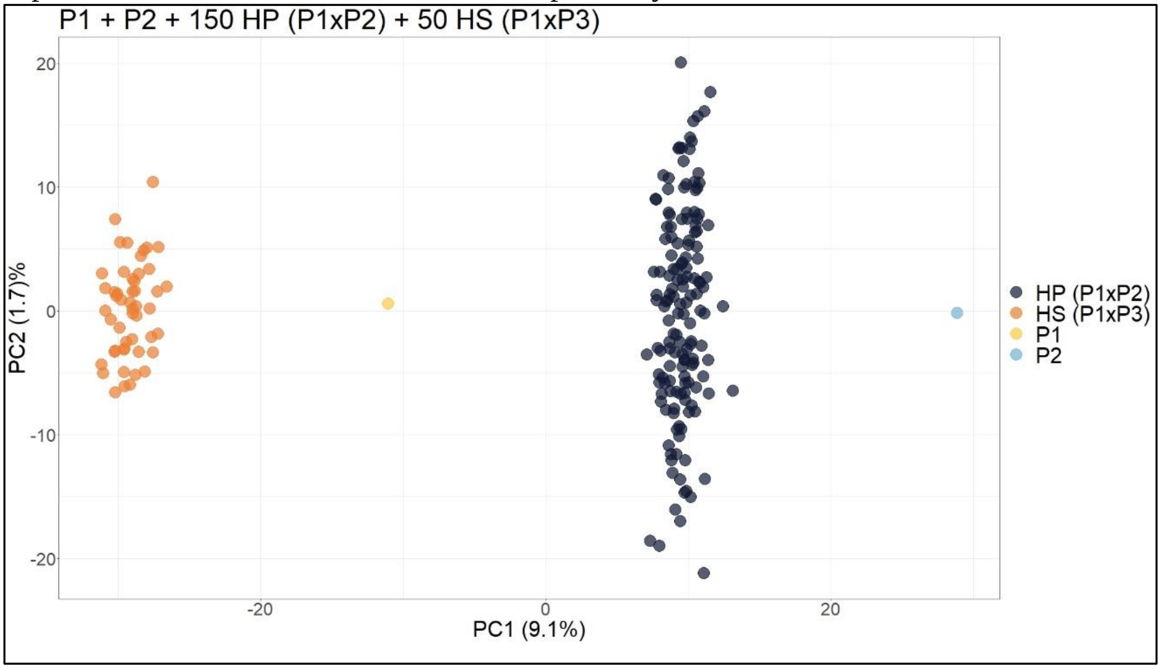


**Supplementary Figure 9**. Principal component analysis scatter plot showing the simulated population with two parents (P1 and P2), 150 hybrids (HP (P1 x P2)) and 50 half-siblings (HS (P1 x P3)). The axes represent the first and second principal components, which explain 9.1% and 1.7% of the variance, respectively.

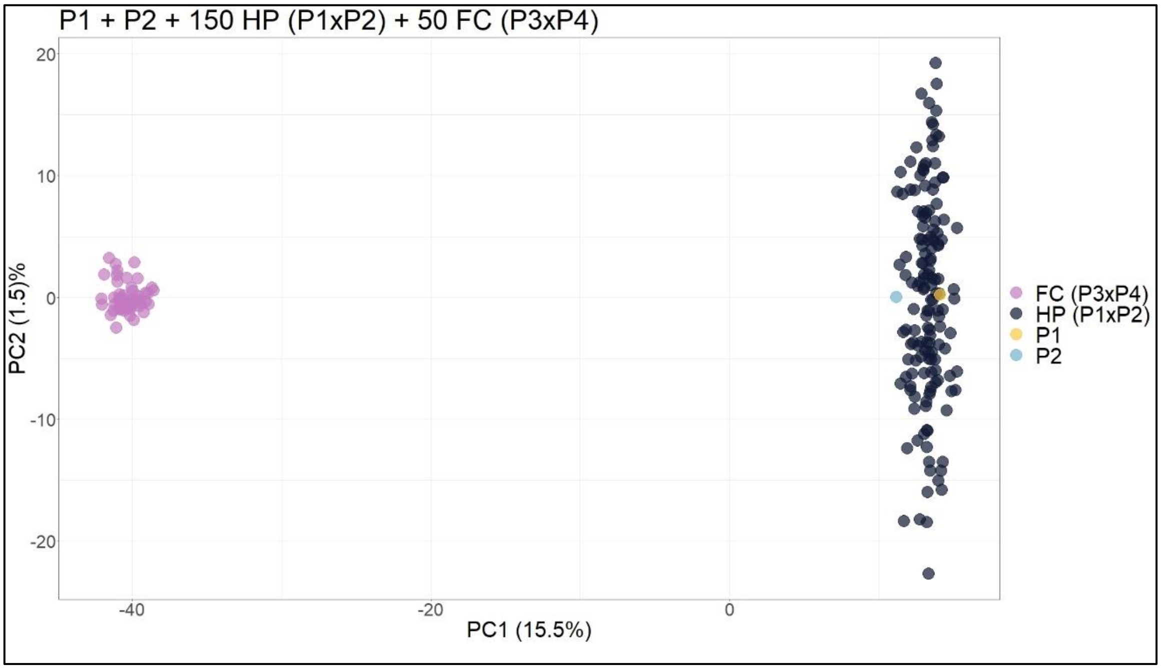


**Supplementary Figure 10**. Principal component analysis scatter plot showing the simulated population with two parents (P1 and P2), 150 hybrids (HP (P1 x P2)) and 50 full contaminants (FC (P3 x P4)). The axes represent the first and second principal components, which explain 15.5% and 1.5% of the variance, respectively.

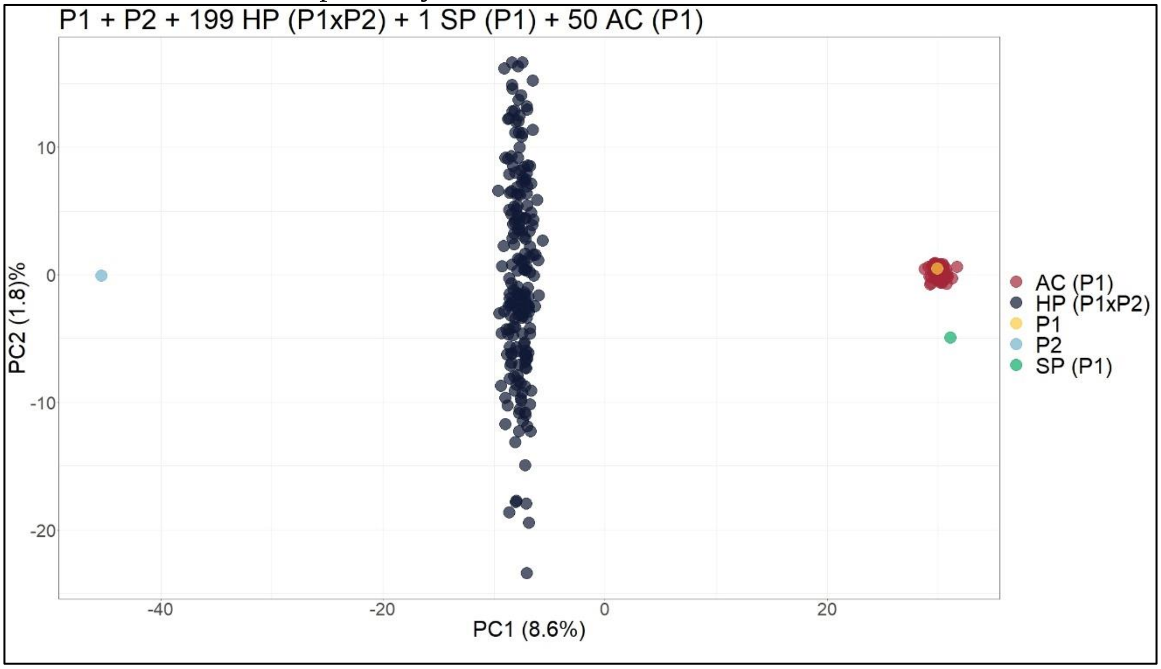


**Supplementary Figure 11**. Principal component analysis scatter plot showing the simulated population with two parents (P1 and P2), 199 hybrids (HP (P1 x P2)), one self-fertilization (SP (P1 x P1)) and 50 apomictic clones (AC (P1)). The axes represent the first and second principal components, which explain 8.6% and 1.8% of the variance, respectively.

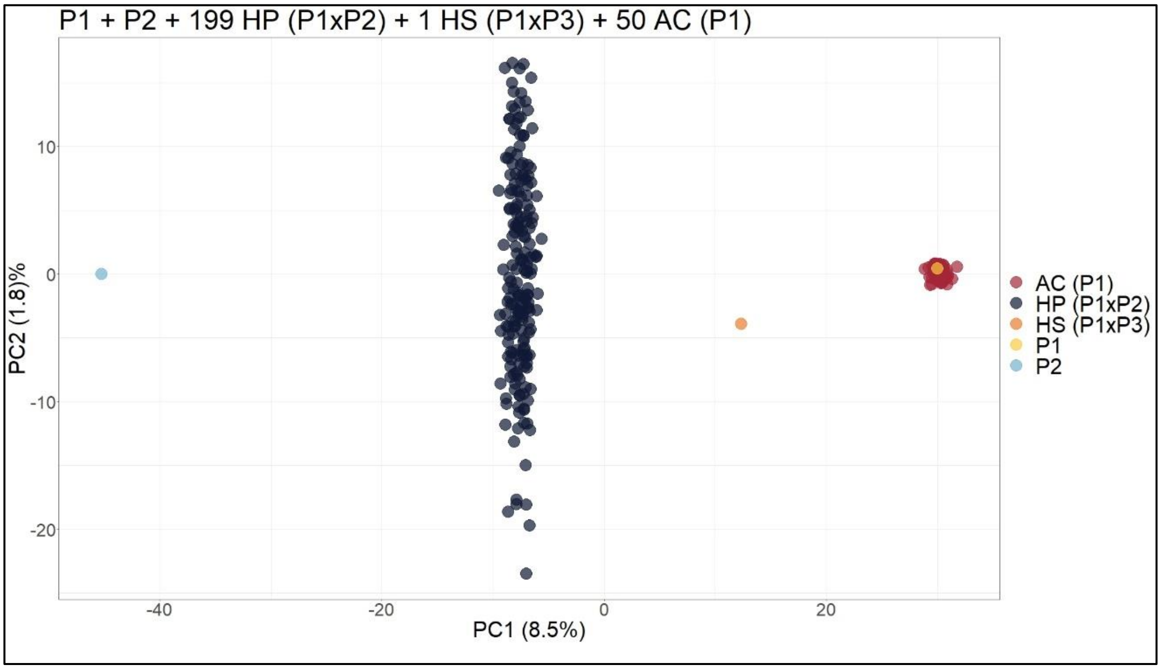


**Supplementary Figure 12**. Principal component analysis scatter plot showing the simulated population with two parents (P1 and P2), 199 HPs (P1 x P2), one half-sibling (HS (P1 x P3)) and 50 apomictic clones (AC (P1)). The axes represent the first and second principal components, which explain 8.5% and 1.8% of the variance, respectively.

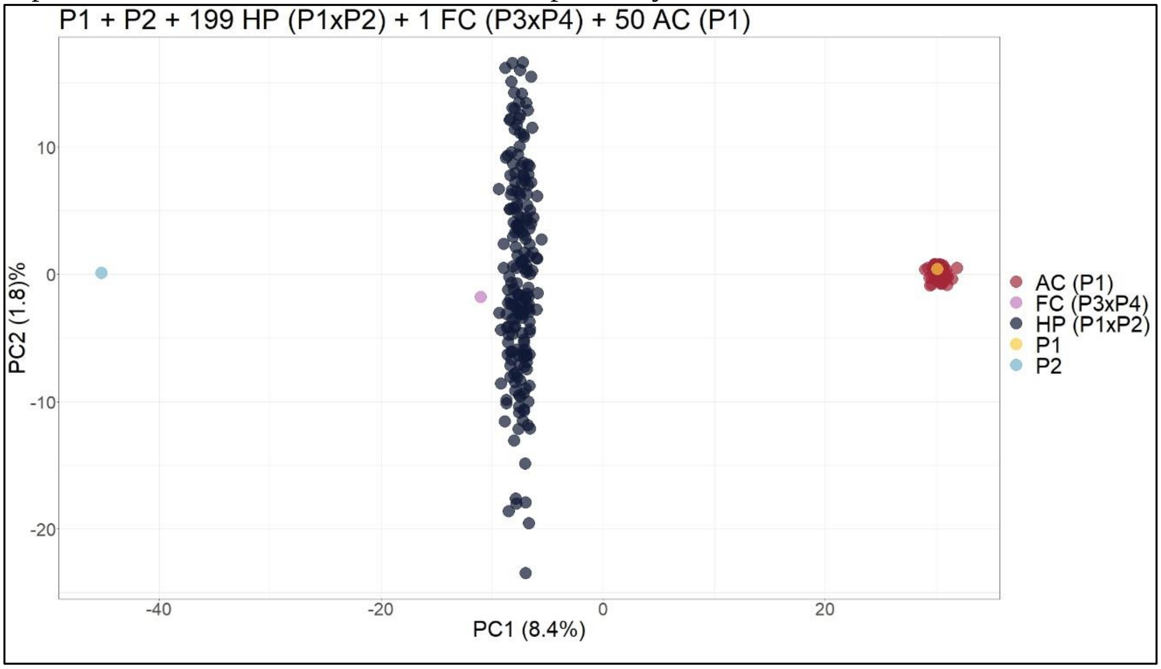


**Supplementary Figure 13**. Principal component analysis scatter plot showing the simulated population with two parents (P1 and P2), 199 HPs (P1 x P2), one full contaminant (FC (P3 x P4)) and 50 apomictic clones (AC (P1)). The axes represent the first and second principal components, which explain 8.4% and 1.8% of the variance, respectively.

##### 2.3 *Megathyrsus maximus*

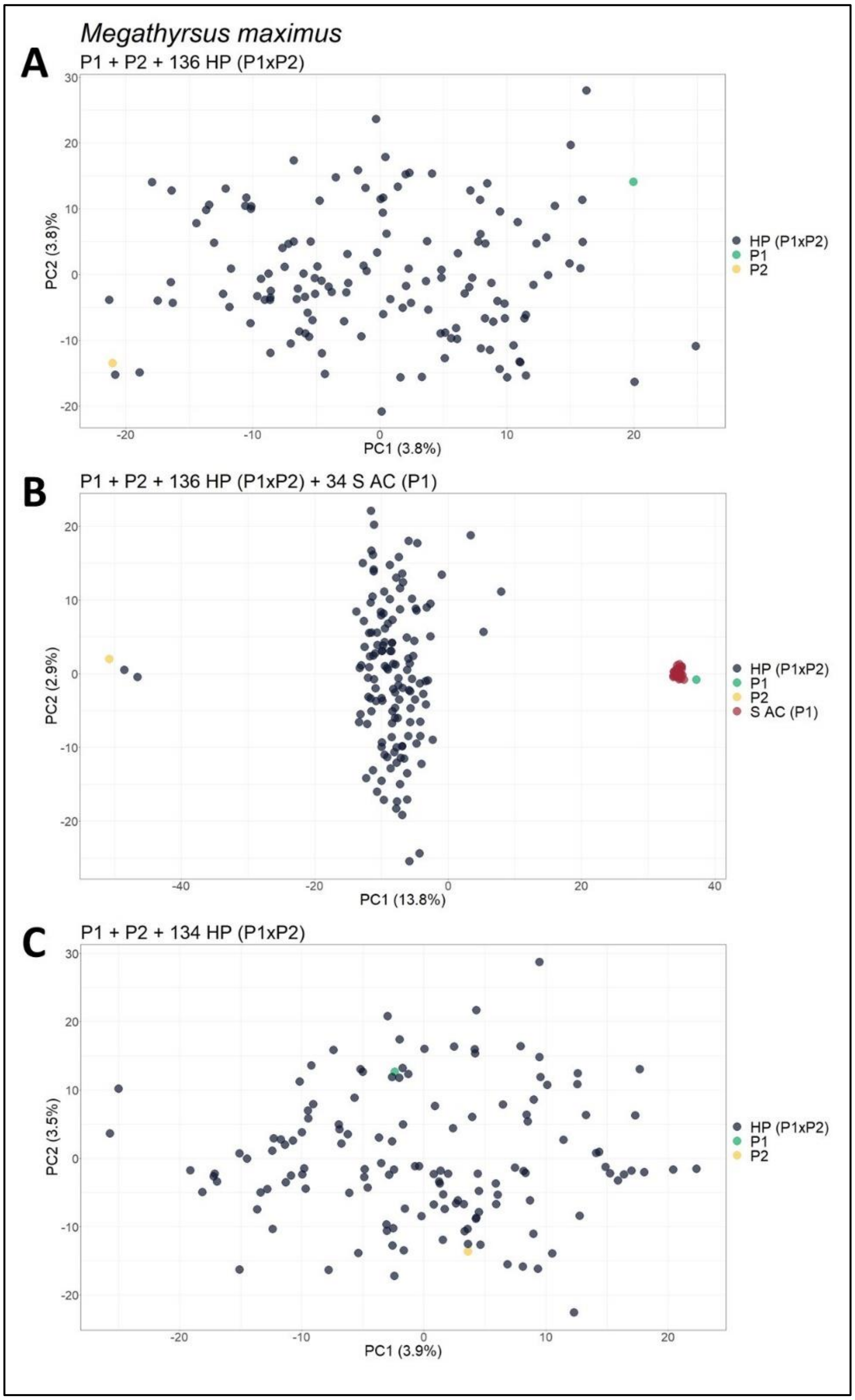


**Supplementary Figure 14**. Principal component analysis (PCA) scatter plots showing the *M. maximus* progeny for the set of SNP markers filtered by a *prop_mis* value of 0.20. (**A**) Original population composed of two parents (P1 and P2) and their progeny of 136 hybrids (HP (P1xP2)); (**B**) Population with 36 simulated apomictic clones (S AC (P1)); (**C**) Population without the 2 apomictic clones (AC) identified. The axes represent the first and second principal components, which explain 3.8% and 3.8% of the variance for (**A**), 13.8% and 2.9% for (**B**), and 3.9% and 3.5% for (**C**), respectively.

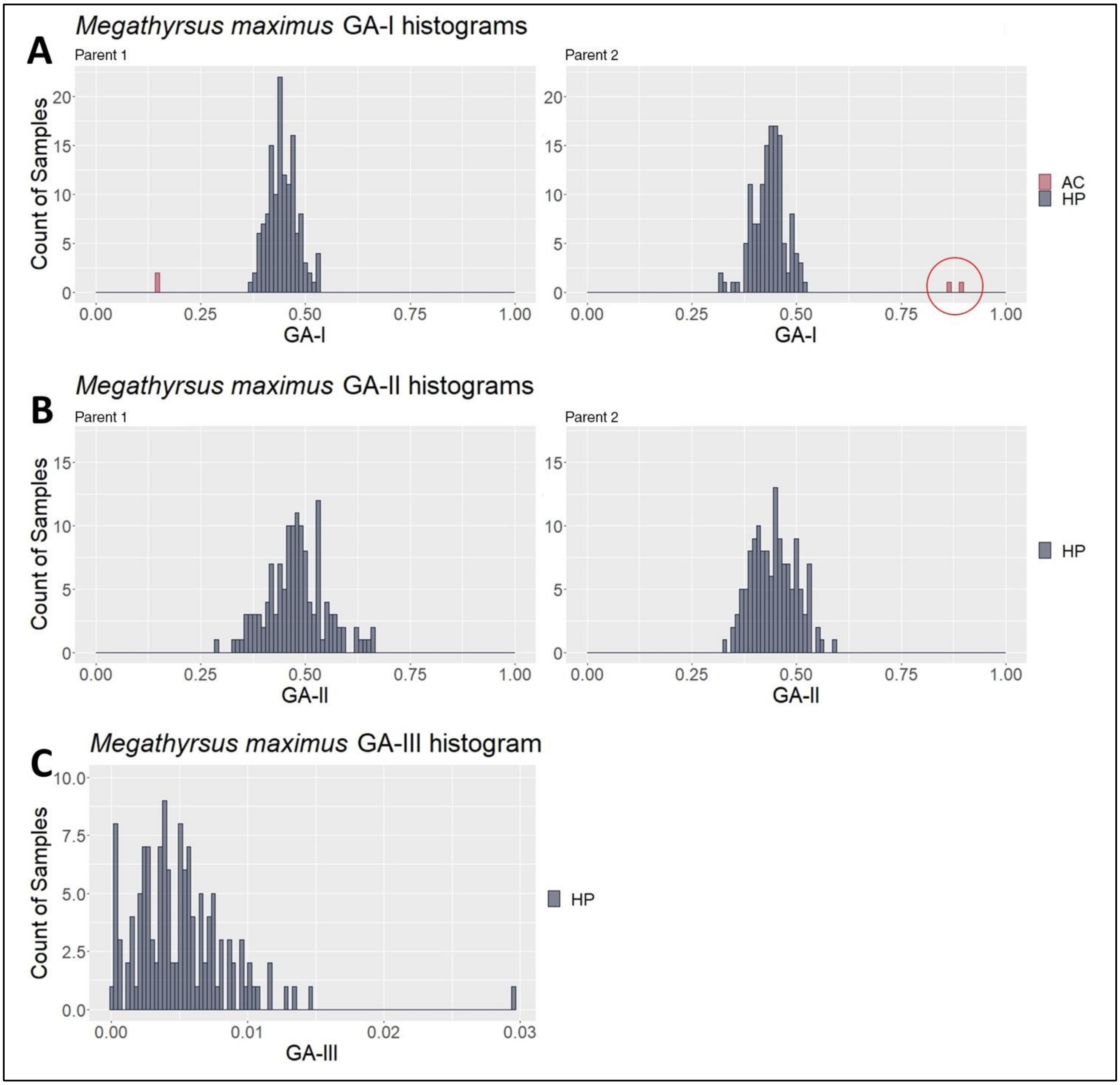


**Supplementary Figure 15**. *Megathyrsus maximus* GA histograms with samples classified by the two clusters obtained in the CA. (**A**), (**B**) and (**C**) show the results for GA-I, GA-II and GA-III, respectively. The red circle highlights the identified AC contaminants in contrast to the hybrid progeny (HP).

#### *2.4 Urochloa humidicola*

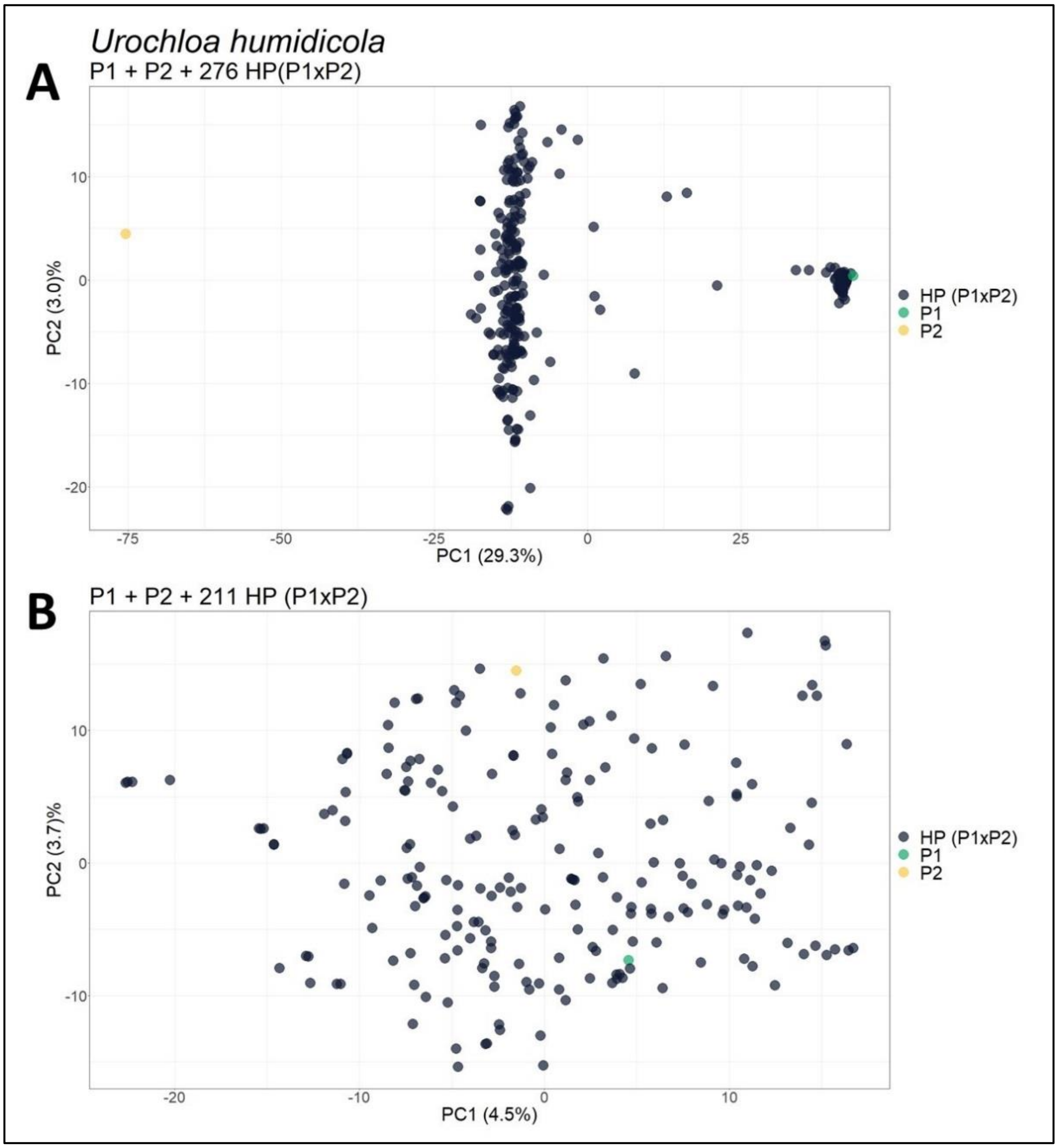


**Supplementary Figure 16**. Principal component analysis (PCA) scatter plots showing the *U. humidicola* progeny for the set of SNP markers filtered by a *prop_mis* value of 0.20. (**A**) Original population composed of two parents (P1 and P2) and their progeny of 276 hybrids (HP (P1xP2)); (**B**) Population without the 65 apomictic clones (AC) identified. The axes represent the first and second principal components, which explain 29.3% and 3.0% of the variance for (**A**) and 4.5% and 3.7% for (**B**), respectively.

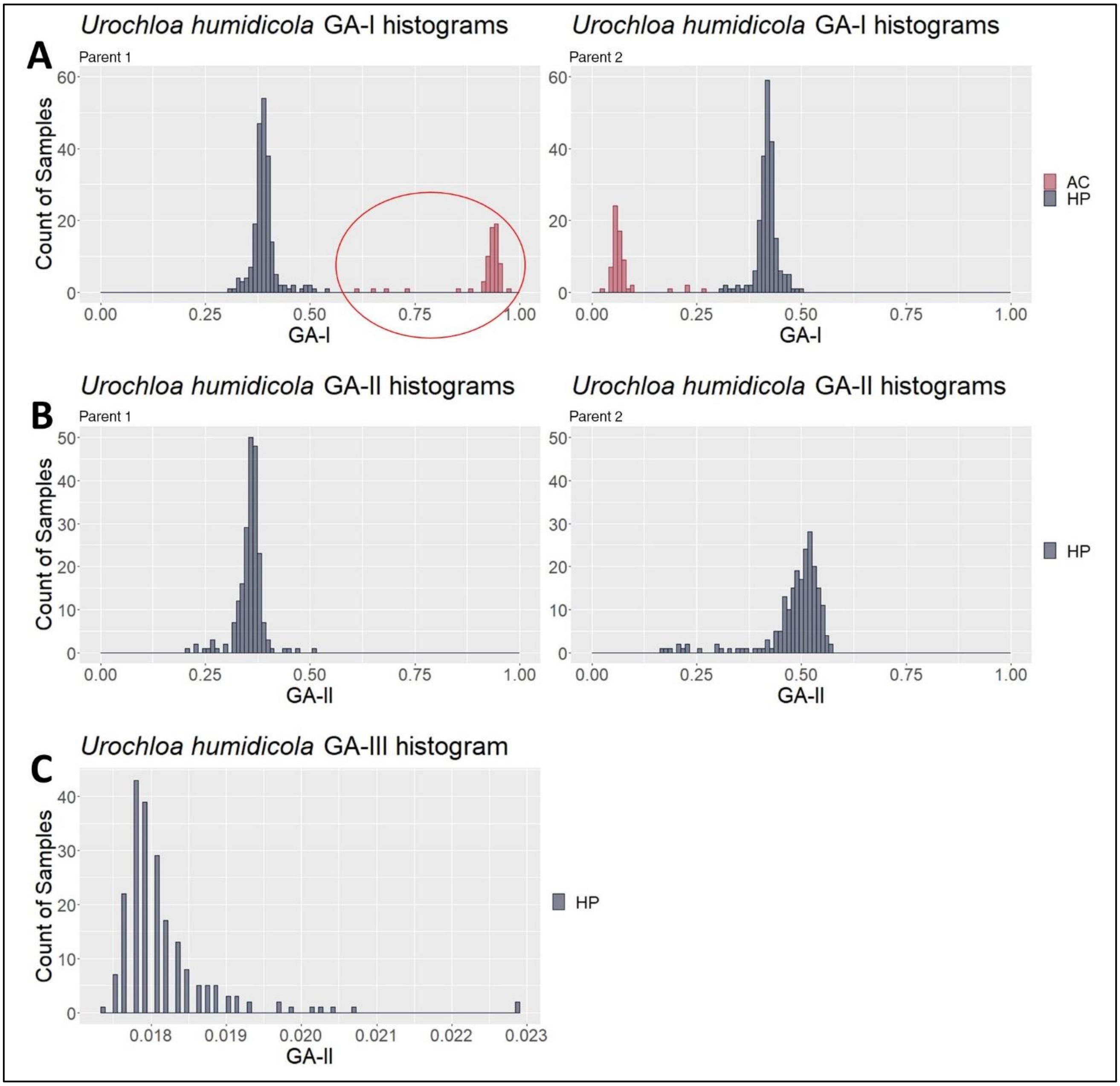


**Supplementary Figure 17**. *Urochloa humidicola* GA histograms with samples classified by the two clusters obtained in the CA. (**A**), (**B**) and (**C**) show the results for GA-I, GA-II and GA-III, respectively. The red circle highlights the identified AC contaminants in contrast to the hybrid progeny (HP).
